# Methylotrophy, alkane-degradation, and pigment production as defining features of the globally distributed yet-uncultured phylum Binatota

**DOI:** 10.1101/2020.09.14.296780

**Authors:** Chelsea L. Murphy, Peter F. Dunfield, Andriy Sheremet, John R. Spear, Ramunas Stepanauskas, Tanja Woyke, Mostafa S. Elshahed, Noha H. Youssef

## Abstract

The recent leveraging of genome-resolved metagenomics has opened a treasure trove of genomes from novel uncultured microbial lineages, yet left many clades undescribed. We here present a global analysis of genomes belonging to the Binatota (UBP10), a globally distributed, yet-uncharacterized bacterial phylum. All orders in the Binatota encoded the capacity for aerobic methylotrophy using methanol, methylamine, sulfomethanes, chloromethanes, and potentially methane as substrates. Methylotrophy in the Binatota was characterized by order-specific substrate degradation preferences, as well as extensive metabolic versatility, i.e. the utilization of diverse sets of genes, pathways and combinations to achieve a specific metabolic goal. The genomes also encoded an arsenal of alkane hydroxylases and monooxygenases, potentially enabling growth on a wide range of alkanes and fatty acids. Pigmentation is inferred from a complete pathway for carotenoids (lycopene, β and γ carotenes, xanthins, chlorobactenes, and spheroidenes) production. Further, the majority of genes involved in bacteriochlorophyll *a*, *c*, and *d* biosynthesis were identified; although absence of key genes and failure to identify a photosynthetic reaction center precludes proposing phototrophic capacities. Analysis of 16S rRNA databases showed Binatota’s preferences to terrestrial and freshwater ecosystems, hydrocarbon-rich habitats, and sponges supporting their suggested potential role in mitigating methanol and methane emissions, alkanes degradation, and nutritional symbiosis with sponges. Our results expand the lists of methylotrophic, aerobic alkane degrading, and pigment-producing lineages. We also highlight the consistent encountering of incomplete biosynthetic pathways and gene shrapnel in microbial genomes, a phenomenon necessitating careful assessment when assigning putative functions based on a set-threshold of pathway completion.

## Introduction

Approaches that directly recover genomes from environmental samples and bypass the hurdle of cultivation (single-cell genomics and genome-resolved metagenomics) have come of age in the last decade. The new availability of environmentally sourced genomes, obtained as SAGs (single amplified genomes) or MAGs (metagenome-assembled genomes) is having a lasting impact on the field of microbial ecology. Distinct, yet often complementary and intertwined, strategies are employed for the analysis of the deluge of obtained genomes. Site- or habitat-specific studies focus on spatiotemporal sampling of a single site or habitat of interest. The obtained genomes are then analyzed to elucidate how resident taxa mediate substrates turnover and elemental cycling, examine microbial interactions on the metabolic and cellular levels, and document how the community responds to natural and anthropogenic changes ^1, 2, 3, 4^. Function-based studies focus on genomes from single or multiple habitats to identify and characterize organisms involved in a specific process, e.g. cellulose degradation ^5^ or sulfate-reduction ^6^. Phylogeny-oriented (phylocentric) studies, on the other hand, focus on characterizing genomes belonging to a specific lineage of interest, with the aim of delineating its pan, core, and dispensable gene repertoire, documenting the defining metabolic capabilities and physiological preferences for the entire lineage and encompassed clades ^7, 8^, understanding the lineage’s ecological distribution and putative roles in various habitats ^4, 9^, and elucidating genomic basis underpinning niche specializing patterns ^10^. The scope of phylocentric studies could range from the analysis of a single genome from a single ecosystem ^11^, to global sampling and *in-silico* analysis efforts ^12, 13^. The feasibility and value of phylocentric strategies have recently been enhanced by the development of a genome-based (phylogenomic) taxonomic outline based on extractable data from MAGs and SAGs providing a solid framework for knowledge building and data communication ^14^, as well as recent efforts for massive, high-throughput binning of genomes from global collections of publicly available metagenomes in GenBank nr and Integrated Microbial Genomes & Microbiomes (IMG/M) databases ^15, 16^.

Candidate phylum UBP10 has originally been described as one of the novel lineages recovered from a massive binning effort that reconstructed thousands of genomes from publicly available metagenomic datasets ^15^. UBP10 has subsequently been named candidate phylum Binatota (henceforth Binatota) in an effort to promote nomenclature for uncultured lineages based on attributes identified in MAGs and SAGs ^17^. The recent generation of 52,515 distinct MAGs binned from over 10,000 metagenomes ^16^ has greatly increased the number of available Binatota genomes. Here, we utilize a phylocentric approach and present a comparative analysis of the putative metabolic and biosynthetic capacities and putative ecological roles of members of the candidate phylum Binatota, as based on sequence data from 108 MAGs. Our study documents aerobic methylotrophy, aerobic alkane degradation, and carotenoid pigmentation as defining traits in the Binatota. We also highlight the presence of incomplete chlorophyll biosynthetic pathways in all genomes, and propose several evolutionary-grounded scenarios that could explain such pattern.

## Results

### Overview

A total of 108 Binatota MAGs with >70% completion and <10% contamination were used for this study, which included 86 medium-quality and 22 high-quality genomes, as based on MIMAG standards ^18^. Binatota genomes clustered into seven orders designated as Bin18 (n=2), Binatales (n=48), HRBin30 (n=7), UBA1149 (n=9), UBA9968 (n=34), UBA12015 (n=1), UTPRO1 (n=7), encompassing12 families, and 24 genera (Figure 1, Table S1). 16S rRNA gene sequences extracted from orders Bin18 and UBA9968 genomes were classified in SILVA (release 138) ^19^ as members of class bacteriap25 in the phylum Myxococcota, order Binatales and order HRBin30 as uncultured phylum RCP2-54, and orders UBA1149 and UTPRO1 as uncultured Desulfobacterota classes (Table S1). RDP II-classification (July 2017 release, accessed July 2020) classified all Binatota sequences as unclassified Deltaproteobacteria (Table S1).

**Figure 1.**
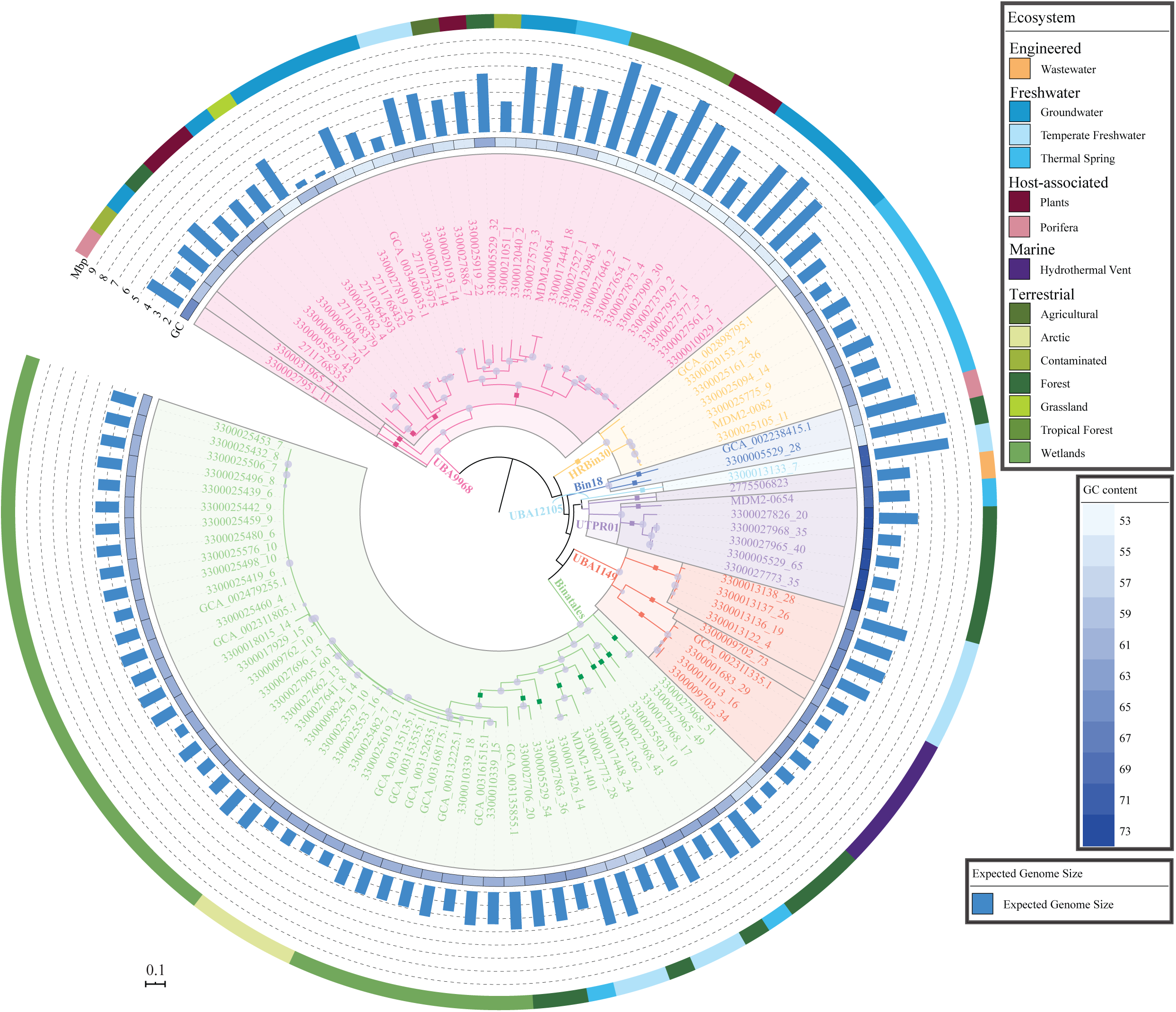
Phylogenomic relationship between analyzed Binatota genomes. The Maximum Likelihood tree was constructed in RAxML from a concatenated alignment of 120 single-copy marker genes. The tree was rooted using *Deferrisoma camini* (GCA_000526155.1) as the outgroup (not shown). Orders are shown as colored wedges: UBA9968, pink; HRBin30, tan; Bin18, blue; UBA12105, cyan; UTPR01, purple; UBA1149, orange; and Binatales, green. Within each order, families are delineated by grey borders, and genera are shown as colored squares on the branches. Bootstrap values are shown as purple bubbles for nodes with *≥*70% support. The tracks around the tree represent (innermost-outermost) GC content (with a heatmap that ranges from 53% (lightest) to 73% (darkest)), expected genome size (bar chart), and classification of the ecosystem from which the genome originated. All genomes analyzed in this study were >70% complete and <10% contaminated. Completion/contamination percentages, and individual genomes assembly size are shown in Tables S2, and S3, respectively.

### Methylotrophy in the Binatota

#### 1. C1 substrates oxidation to formaldehyde

##### Methanol

With the exception of HRBin30, all orders encoded at least one type of methanol dehydrogenase (Figure 2a). Three distinct types of methanol dehydrogenases were identified (Figure 2a, b): 1. The NAD(P)-binding MDO/MNO-type methanol dehydrogenase (*mno*), typically associated with Gram-positive methylotrophic bacteria (Actinobacteria and *Bacillus methanolicus*) ^20^, was the only type of methanol dehydrogenase identified in orders UBA9968, UBA12105, and UTPR01 (Figure 2a, Extended data 1), as well as some UBA1149 and Binatales genomes. 2. The MDH2-type methanol dehydrogenase, previously discovered in members of the Burkholderiales and Rhodocyclales ^21^, was encountered in the majority of order UBA1149 genomes and in two Binatales genomes, and 3. The lanthanide-dependent pyrroloquinoline quinone (PQQ) methanol dehydrogenase XoxF-type was encountered in nine genomes from the orders Bin18, and Binatales, together with the accessory XoxG (c-type cytochrome) and XoxJ (periplasmic binding) proteins (Figure 2a). All later genomes also encoded PQQ biosynthesis. Surprisingly, none of the genomes encoded the MxaF1-type (MDH1) methanol dehydrogenase, typically encountered in model methylotrophs ^22^.

**Figure 2.**
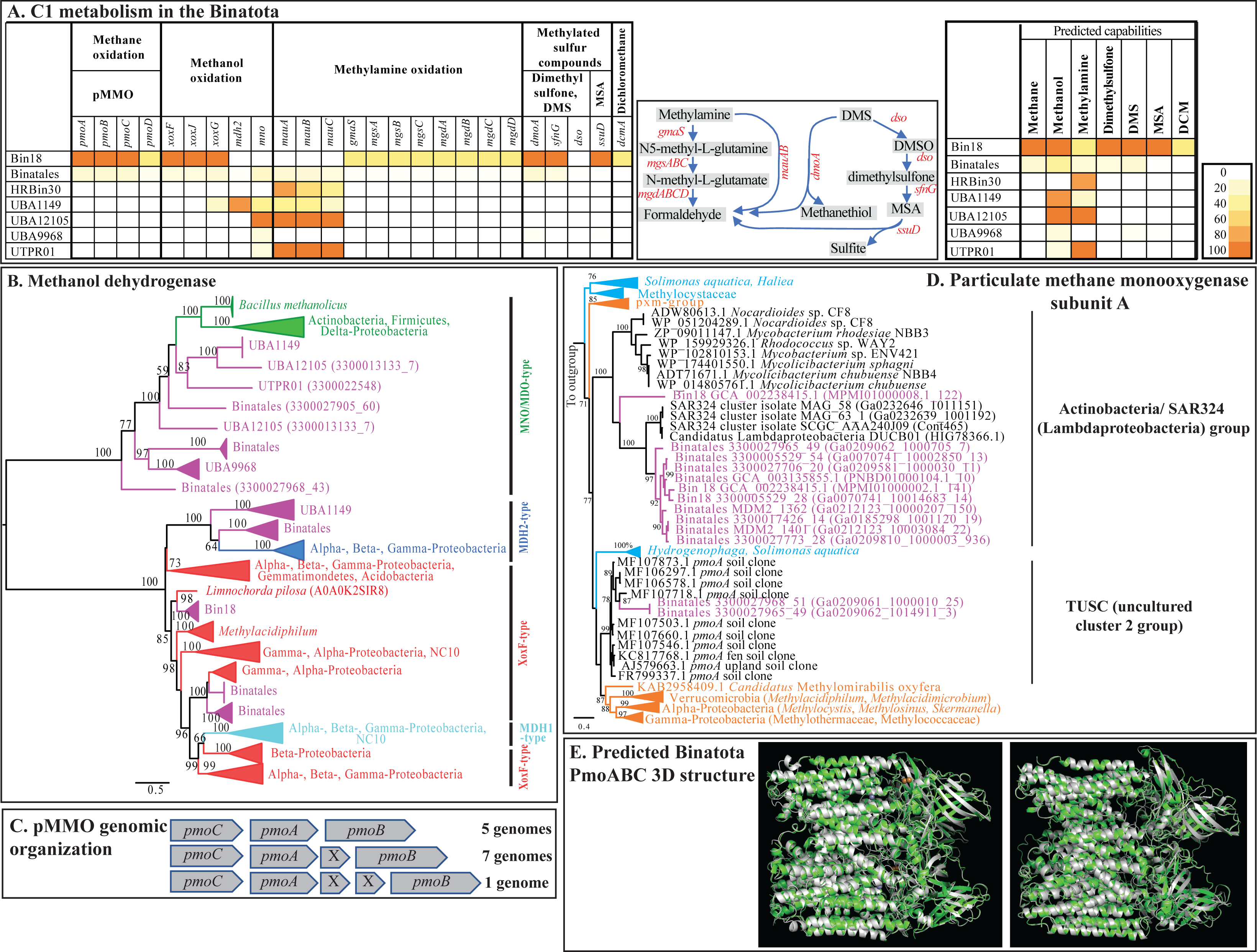
C1 substrate degradation capacities in the Binatota. (A) Heatmap of the distribution of various C1 oxidation genes in Binatota genomes from different orders. The heatmap colors (as explained in the key) correspond to the percentage of genomes in each order encoding a homologue of the gene in the column header. Pathways involving more than one gene for methylamine and methylated sulfur compounds degradation are shown next to the heatmap. To the right, the per-order predicted C1 oxidation capacity is shown as a heatmap with the colors corresponding to the percentage of genomes in each order where the full degradation pathway was detected for the substrate in the column header. These include *pmoABC* for methane, *xoxFJG, mdh2*, and/or *mno* for methanol, *mau* and/or indirect glutamate pathway for methylamine, *sfnG* and *ssuD* for dimethylsulfone, *dso*, *sfnG* and *ssuD*, or *dmoA* for dimethylsulfide (DMS), *ssuD* for methane sulfonic acid (MSA), and *dcmA* for dichloromethane (DCM). pMMO: particulate methane monooxygenase with *pmoA*, *pmoB*, *pmoC, pmoD* denoting subunits A, B, C, and D; XoxF-type (*xoxF*, *xoxJ*, *xoxG*), MDH2-type (*mdh2*), and MNO/MDO-type (*mno*) methanol dehydrogenases; direct oxidation methylamine dehydrogenase (*mauABC*), indirect glutamate pathway (*gmaS*: *γ*-glutamylmethylamide synthase; *mgsABC*: *N*-methyl-L-glutamate synthase; methylglutamate dehydrogenase *mgdABCD*); dimethylsulfide (DMS) monooxygenase (*dmoA*), dimethyl sulfone monooxygenase (*sfnG*), dimethylsulfide monooxygenase (*dso*), alkane sulfonic acid monooxygenase (*ssuD*); and dichloromethane dehalogenase (*dcmA*)). (B) Maximum likelihood phylogenetic tree highlighting the relationship between Binatota methanol dehydrogenases in relation to other methylotrophic taxa. Bootstrap support (from 100 bootstraps) is shown for branches with >50%. (C) Organization of pMMO genes in Binatota genomes, and the number of genomes where each organization was observed. x: Hypothetical protein (D) Maximum likelihood tree highlighting the relationship between Binatota *pmoA* genes to methanotrophic taxa and environmental amplicons. Bootstrap support (100 bootstraps) is shown for branches with >50% support. Sequences from Binatota genomes (shown as Order followed by Bin name then pmoA protein ID in parentheses) are in magenta and fall into two clusters; Actinobacteria/SAR324 cluster, and TUSC uncultured cluster 2. Clusters from previously studied pMMOs known to reduce methane are in orange, while those known to reduce short chain alkanes but not methane are in cyan (collective data from ^28, 29, 30, 131, 132, 133^). The tree was rooted using the amoA sequence of *Candidatus* Nitrosarchaeum limnium SFB1 (EGG41084.1) as an outgroup. (E) Predicted particulate methane monooxygenase (PmoABC) 3D structure (grey) from a Cluster 2 TUSC-affiliated Binatota genome (Genome 3300027968_51, left), and an Actinobacteria/SAR324-affiliated Binatota genome (Genome GCA_002238415.1, right) both superimposed on pMMO from the model methanotroph *Methylococcus capsulatus* str. Bath (pdb: 3RGB) (green) with a global model quality estimate of 0.7, and 0.62, respectively, and a quaternary structure quality score of 0.57, and 0.55, respectively.

##### Methylamine

All Binatota orders except UBA9968 encoded methylamine degradation capacity. The direct periplasmic route (methylamine dehydrogenase; *mau*) was more common, with *mauA* and *mauB* enzyme subunits encoded in the Binatales, HRBin30, UBA1149, UBA12105, and UTPR01 (Figure 2a, Extended data 1). Amicyanin (encoded by *mauC*) is the most probable electron acceptor for methylamine dehydrogenase ^22^ (Figure 2a). On the other hand, one Bin18 genome, and two Binatales genomes (that also encode the *mau* cluster) encoded the full complement of genes for methylamine oxidation via the indirect glutamate pathway (Figure 2a, Extended data 1).

##### Methylated sulfur compounds

Binatota genomes encoded several enzymes involved in the degradation of dimethyl sulfone, methane sulfonic acid (MSA), and dimethyl sulfide (DMS). Nine genomes (two Bin18, and 7 Binatales) encoded dimethyl sulfone monooxygenase (*sfnG*) involved in the degradation of dimethyl sulfone to MSA with the concomitant release of formaldehyde. Three of these nine genomes also encoded alkane sulfonic acid monooxygenase (*ssuD*), which will further degrade the MSA to formaldehyde and sulfite. Degradation of DMS via DMS monooxygenase (*dmoA*) to formaldehyde and sulfide was encountered in 13 genomes (2 Bin18, 9 Binatales, and 2 UBA9968). Further, one Binatales genome encoded the *dso* system (EC: 1.14.13.245) for DMS oxidation to dimethyl sulfone, which could be further degraded to MSA as explained above (Figure 2a, Extended data 1).

##### Dihalogenated methane

One Bin18 genome encoded the specific dehalogenase/ glutathione S-transferase (*dcmA*) capable of converting dichloromethane to formaldehyde.

##### Methane

Genes encoding particulate methane monooxygenase (pMMO) were identified in orders Bin18 (2/2 genomes) and Binatales (9/48 genomes) (Figure 2a, Extended data 1), while genes encoding soluble methane monooxygenase (sMMO) were not found. A single copy of all three pMMO subunits (A, B, and C) was encountered in 9 of the 11 genomes, while two copies were identified in two genomes. pMMO subunit genes (A, B, and C) occurred as a contiguous unit in all genomes, with a CAB (5 genomes), and/or CAxB or CAxxB (8 genomes, where x is a hypothetical protein) organization, similar to the pMMO operon structure in methanotrophic Proteobacteria, Verrucomicrobia, and *Candidatus* Methylomirabilis (NC10) ^23, 24, 25, 26^ (Figure 2c)). In addition, five of the above eleven genomes also encoded a *pmoD* subunit, recently suggested to be involved in facilitating the enzyme complex assembly, and/or in electron transfer to the enzyme’s active site ^27, 28^. Phylogenetic analysis of Binatota *pmoA* sequences revealed their affiliation with two distinct clades: the yet-uncultured Cluster 2 TUSC (<u>T</u>ropical <u>U</u>pland <u>S</u>oil <u>C</u>luster) methanotrophs ^29^ (2 Binatales genomes), and a clade encompassing *pmoA* sequences from Actinobacteria (*Nocardioides* sp. strain CF8, *Mycolicibacterium*, and *Rhodococcus*) and SAR324 (*Candidatus* Lambdaproteobacteria) ^30, 31^ (Figure 2d). Previous studies have linked Cluster 2 TUSC pMMO-harboring organisms to methane oxidation based on selective enrichment on methane in microcosms derived from Lake Washington sediments ^32^. All Binatota genomes encoding TUSC-affiliated pMMO, also encoded genes for downstream methanol and formaldehyde oxidation as well as formaldehyde assimilation (see below), providing further evidence for their putative involvement in methane oxidation. On the other hand, studies on *Nocardioides* sp. strain CF8 demonstrated its capacity to oxidize short chain (C2-C4) hydrocarbons, but not methane, via its pMMO, and its genome lacked methanol dehydrogenase homologues ^33^. Such data favor a putative short chain hydrocarbon degradation function for organisms encoding this type of pMMO, although we note that five out of the nine Binatota genomes encoding SAR324/ Actinobacteria-affiliated *pmoA* sequences also encoded at least one methanol dehydrogenase homologue. Modeling pMMO subunits from both TUSC-type and Actinobacteria/SAR324-type Binatota genomes using *Methylococcus capsulatus* (Bath) 3D model (PDB ID: 3rbg) revealed a heterotrimeric structure (*α*_3_*β*_3_*γ*_3_) with the 7, 2, and 5 alpha helices of the PmoA, PmoB, and PmoC subunits, respectively, as well as the beta sheets characteristic of PmoA, and PmoB subunits (Figure 2e). Modeling also predicted binding pockets of the dinuclear Cu ions and Zn ligands (Figure 2e).

#### 2. Formaldehyde oxidation to CO_2_

Three different routes for formaldehyde oxidation to formate were identified (Figure 3). First, the Actinobacteria specific thiol-dependent formaldehyde dehydrogenase (*fadh/mscR*) (EC: 1.1.1.306) was, surprisingly, detected in the majority (96 out of 108) of genomes (Figure 3a, Extended data 1). The enzyme requires a specific thiol (mycothiol ^34^), the biosynthesis of which (encoded by *mshABC* gene cluster) was also encoded in Binatota genomes (Figure 3a). Second, the tetrahydrofolate (H_4_F)-linked pathway comprising the genes *folD* (encoding bifunctional methylene-H_4_F dehydrogenase and methenyl-H_4_F cyclohydrolase) and either *ftfL* (the reversible formyl-H_4_F ligase) or *purU* (the irreversible formyl-H_4_F hydrolase) was also widespread (98/108 genomes). Finally, 40 genomes (Bin18, Binatales, HRBin30, and UTPR01) also encoded the single gene/enzyme NAD-linked glutathione-independent formaldehyde dehydrogenase *fdhA*. Surprisingly, no evidence of the most common formaldehyde oxidation pathway (tetrahydromethanopterin (H_4_MPT)-linked) was detected in any of the Binatota genomes. The NAD- and glutathione-dependent formaldehyde oxidation pathway was found incomplete: while homologs of formaldehyde dehydrogenase (*frmA*) were detected in almost all Binatota genomes, S-formylglutathione hydrolase (*frmB*) were absent. Following formaldehyde oxidation to formate, formate is subsequently oxidized to CO_2_ by one of many formate dehydrogenases. The majority of Binatota genomes (103/108) encoded at least one copy of the NAD-dependent formate dehydrogenase (EC: 1.17.1.9) (Figure 3a, Extended data 1).

**Figure 3.**
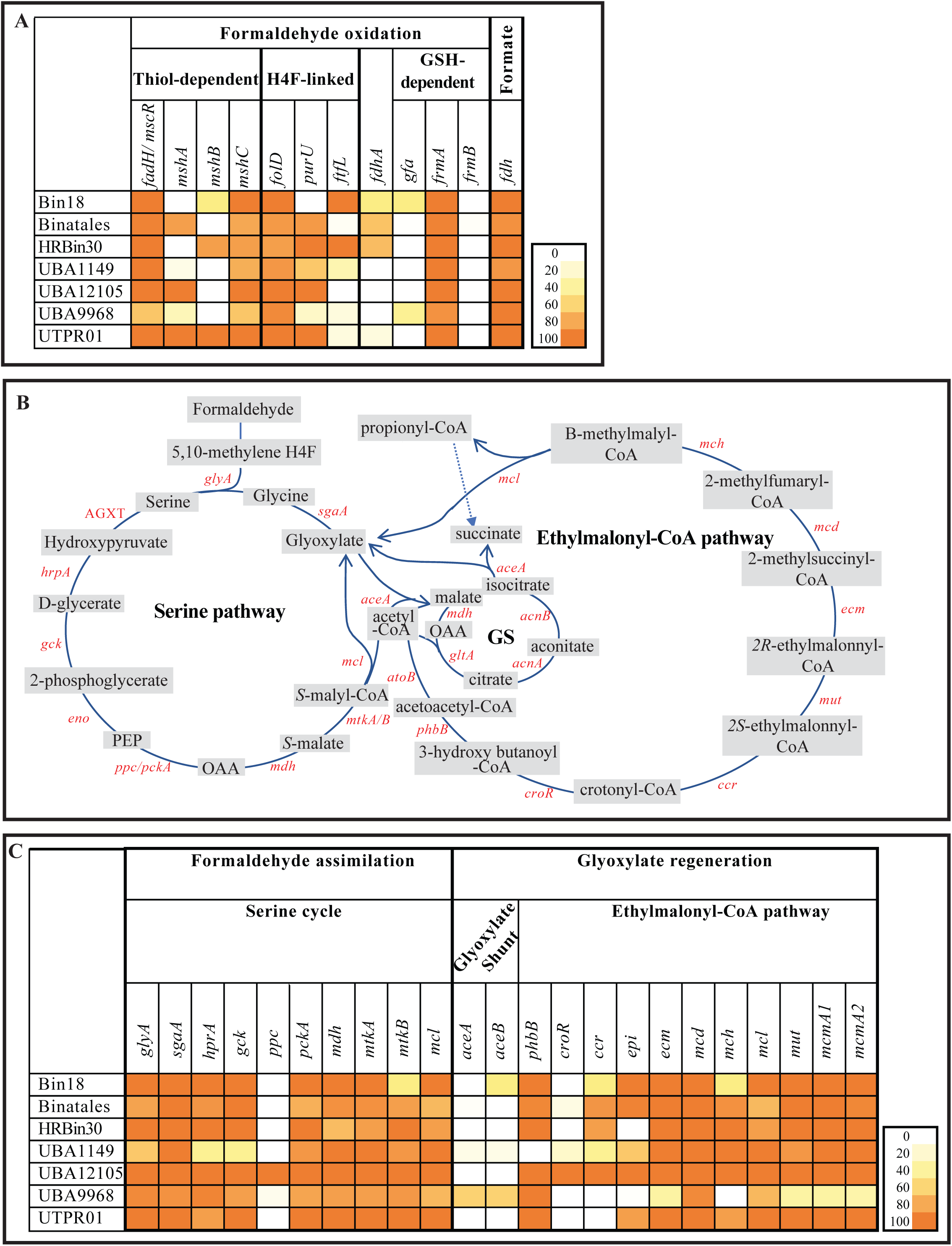
Formaldehyde oxidation and assimilation capabilities encoded by Binatota genomes. (A) Heatmap of the distribution of formaldehyde oxidation genes in Binatota genomes from different orders. The heatmap colors (as explained in the key) correspond to the percentage of genomes in each order encoding a homologue of the gene in the column header. Shown are the different routes of formaldehyde oxidation, including the (myco)thiol-dependent formaldehyde dehydrogenase *fadH/mscR* (along with mycothiol biosynthesis genes (*mshABC*)), the H_4_F-linked pathway (comprising the genes bifunctional methylene-H_4_F dehydrogenase and methenyl-H_4_F cyclohydrolase (*folD*), reversible formyl-H_4_F ligase (*ftfL*), irreversible formyl-H_4_F hydrolase (*purU*)), the glutathione-independent formaldehyde dehydrogenase (*fdhA*), and the glutathione-dependent formaldehyde (comprising the S-(hydroxymethyl)glutathione synthase (*gfa*), NAD- and glutathione-dependent formaldehyde dehydrogenase (*frmA*), S-formylglutathione hydrolase (*frmB*)). Also shown is the distribution of the NAD-dependent formate dehydrogenase (EC: 1.17.1.9) (*fdh*) for formate oxidation. (B) Overview of the pathways for formaldehyde assimilation via the serine cycle (left), and glyoxylate regeneration via the ethylmalonyl-CoA pathway and the glyoxylate shunt (GS) (right). Names of enzymes are shown in red and their distribution in the Binatota genomes from different orders is shown in the heatmap in (C). *glyA*, glycine hydroxymethyltransferase [EC:2.1.2.1]; *sgaA*; serine-glyoxylate transaminase [EC:2.6.1.45]; *hprA*, glycerate dehydrogenase [EC:1.1.1.29]; *gck*, glycerate 2-kinase [EC:2.7.1.165]; *ppc*, phosphoenolpyruvate carboxylase [EC:4.1.1.31]; *pckA*, phosphoenolpyruvate carboxykinase; *mdh*, malate dehydrogenase [EC:1.1.1.37]; *mtkA/B*, malate-CoA ligase [EC:6.2.1.9]; *mcl*, malyl-CoA/(S)-citramalyl-CoA lyase [EC:4.1.3.24 4.1.3.25]; *aceA*, isocitrate lyase [EC:4.1.3.1]; *aceB*, malate synthase [EC:2.3.3.9]; *phbB*, acetoacetyl-CoA reductase [EC:1.1.1.36]; *croR*, 3-hydroxybutyryl-CoA dehydratase [EC:4.2.1.55]; *ccr*, crotonyl-CoA carboxylase/reductase [EC:1.3.1.85]; *epi*, methylmalonyl-CoA/ethylmalonyl-CoA epimerase [EC:5.1.99.1]; *ecm*, ethylmalonyl-CoA mutase [EC:5.4.99.63]; *mcd*, (2S)-methylsuccinyl-CoA dehydrogenase [EC:1.3.8.12]; *mch*, 2-methylfumaryl-CoA hydratase [EC:4.2.1.148]; *mut*, methylmalonyl-CoA mutase [EC:5.4.99.2]; *mcmA1/A2*, methylmalonyl-CoA mutase [EC:5.4.99.2]. Abbreviations: PEP, phosphoenol pyruvate; OAA, oxaloacetate.

#### 3. Formaldehyde assimilation

Two pathways for formaldehyde assimilation by methylotrophs have been described: the serine cycle, which assimilates 2 formaldehyde molecules and 1 CO_2_ molecule, and the ribulose monophosphate cycle (RuMP), which assimilates 3 formaldehyde and no CO_2_ molecules. In addition, some methylotrophs assimilate carbon at the level of CO_2_ via the Calvin Benson Bassham (CBB) cycle ^22^. Homologs encoding the RuMP cycle-specific enzymes were missing from all Binatota genomes, and only three genomes belonging to the Binatales order encoded the CBB cycle enzymes phosphoribulokinase and rubisCO. On the other hand, genes encoding enzymes of the serine cycle (Figure 3b) were identified in all genomes (Figure 3c, Extended data 1), with the key enzymes that synthesize and cleave malyl-CoA (*mtkA/B* [EC 6.2.1.9] malate-CoA ligase, and *mcl* [EC 4.1.3.24] malyl-CoA lyase, respectively) encountered in 98, and 86 Binatota genomes, respectively (Figure 3c, Extended data 1). The entry point of CO_2_ to the serine cycle is the phosphoenolpyruvate (PEP) carboxylase (*ppc*) step catalyzing the carboxylation of PEP to oxaloacetate (Figure 3b). Homologues of *ppc* were missing from most Binatota genomes. Instead, all genomes encoded PEP carboxykinase (*pckA*) that replaces *ppc* function as shown in methylotrophic mycobacteria ^35^ (Figure 3b-c, Extended data 1).

During the serine cycle, regeneration of glyoxylate from acetyl-CoA is needed to restore glycine and close the cycle. Glyoxylate regeneration can be realized either through the classic glyoxylate shunt ^36^, or the ethylmalonyl-CoA pathway (EMCP) ^37^ (Figure 3b). All Binatota genomes exhibited the capacity for glyoxylate regeneration, but the pathway employed appears to be order-specific. Genes encoding all EMCP pathway enzymes were identified in genomes belonging to the orders Bin18, Binatales, HRBin30, UBA1149, and UBA12105 (Figure 3c, Extended data 1), including the two EMCP-specific enzymes ethylmalonyl-CoA mutase (*ecm*) and crotonyl-CoA reductase/carboxylase (*ccr*). On the other hand, order UBA9968 genomes lacked EMCP-specific enzymes but encoded the classic glyoxylate shunt enzymes isocitrate lyase (*aceA*) and malate synthase (*aceB*) (Figure 3c, Extended data 1).

### Alkane degradation

Besides methylotrophy and methanotrophy, Binatota genomes exhibited extensive short-, medium-, and long-chain alkanes degradation capabilities. In addition to the putative capacity of Actinobacteria/SAR324-affiliated pMMO to oxidize C_1_-C_5_ alkanes, and C_1_-C_4_ alkenes as described above, some Binatota genomes encoded propane-2-monoxygenase (*prmABC*), an enzyme mediating propane hydroxylation in the 2-position yielding isopropanol. Several genomes, also encoded medium chain-specific alkane hydroxylases, e.g. homologues of the nonheme iron *alkB* ^38^ and Cyp153-class alkane hydroxylases ^39^. The genomes also encoded multiple long-chain specific alkane monooxygenase, e.g. *ladA* homologues (EC:1.14.14.28) ^40^ (Figure 4a, Extended data 1). Finally, Binatota genomes encoded the capacity to metabolize medium-chain haloalkane substrates. All genomes encoded *dhaA* (haloalkane dehalogenases [EC:3.8.1.5]) known to have a broad substrate specificity for medium chain length (C3 to C10) mono-, and dihaloalkanes, resulting in the production of their corresponding primary alcohol, and haloalcohols, respectively ^41^ (Figure 4a, Extended data 1).

**Figure 4.**
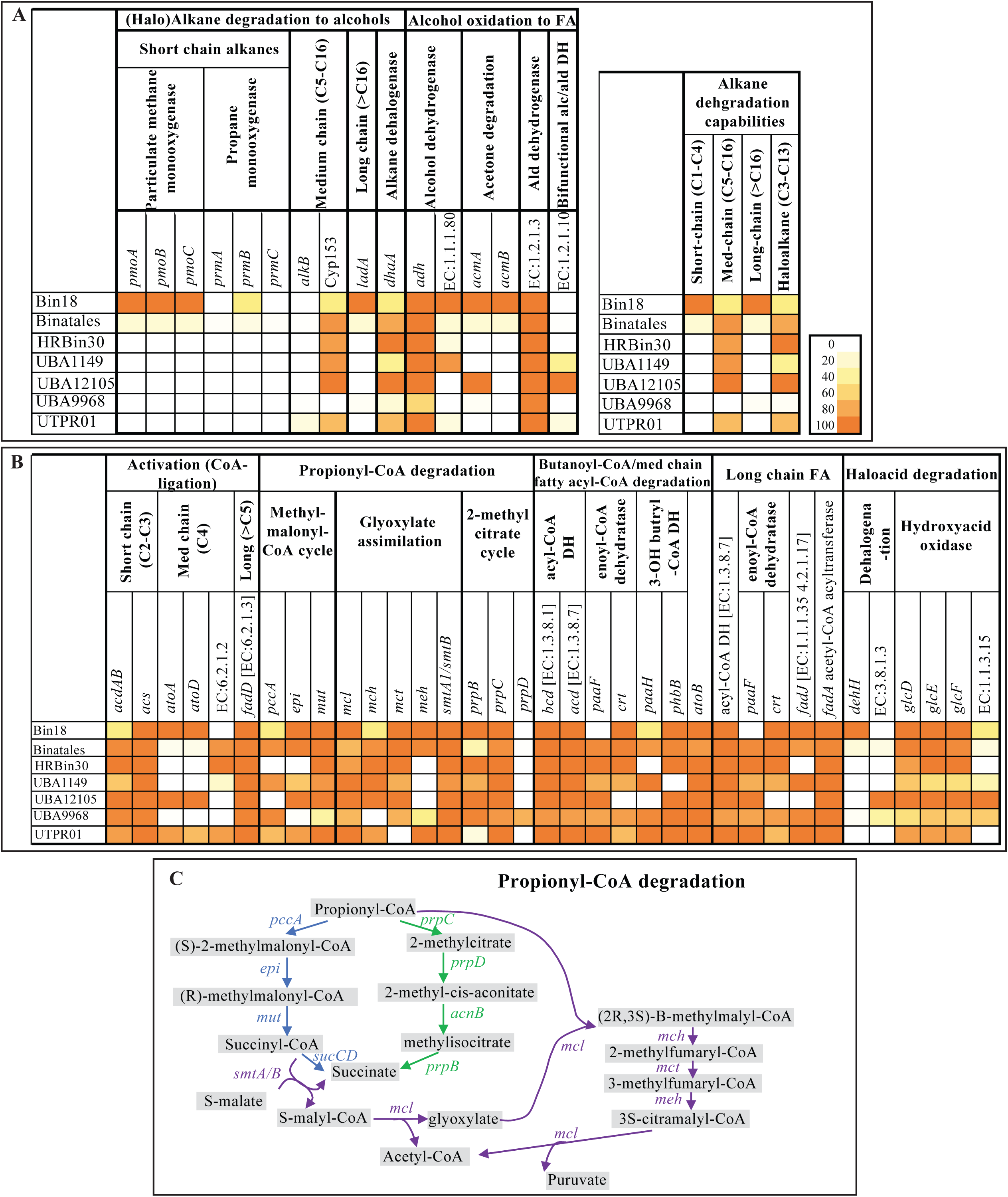
Alkane, and fatty acid degradation capabilities encoded in Binatota genomes. (A) Heatmap of the distribution of (halo)alkane degradation to alcohol. The heatmap colors (as explained in the key) correspond to the percentage of genomes in each order encoding a homologue of the gene in the column header. The per-order predicted alkane degradation capacity is shown to the right as a heatmap with the colors corresponding to the percentage of genomes in each order where the full degradation pathway was detected for the substrate in the column header. These include *pmoABC* and/or *prmABC* for short-chain alkanes, *alkB*, or cyp153 for medium-chain alkanes, *ladA* for long-chain alkanes, and *dhaA* for haloalkanes. (B) Heatmap of the distribution of various chain-length fatty acid and haloacid degradation genes in Binatota genomes. The heatmap colors (as explained in the key) correspond to the percentage of genomes in each order encoding a homologue of the gene in the column header. (C) Propionyl-CoA degradation pathways encoded by the Binatota genomes. The methylmalonyl CoA (MMCoA) pathway is shown in blue, while the 2-methylcitrate pathway is shown in green. In some genomes, the MMCoA pathway seems to be functional but with a slight modification (shown in purple) that includes glyoxylate assimilation and regeneration. *pmoABC*, particulate methane monooxygenase with denoting subunits A, B, and C; *prmABC*, propane 2-monooxygenase [EC:1.14.13.227]; *alkB*, alkane 1-monooxygenase [EC:1.14.15.3]; cyp153, Cytochrome P450 alkane hydroxylase [EC 1.14.15.1]; *ladA*, long-chain alkane monooxygenase [EC:1.14.14.28]; *dhaA*, haloalkane dehalogenase [EC:3.8.1.5]; *adh*, alcohol dehydrogenase [EC:1.1.1.1]; EC:1.1.1.80, isopropanol dehydrogenase (NADP+) [EC:1.1.1.80]; *acmA*, acetone monooxygenase (methyl acetate-forming) [EC:1.14.13.226]; *acmB*, methyl acetate hydrolase [EC:3.1.1.114]; EC:1.2.1.3, aldehyde dehydrogenase (NAD+) [EC:1.2.1.3]; E1.2.1.10, acetaldehyde dehydrogenase (acetylating) [EC:1.2.1.10]; *acdAB*, acetate---CoA ligase (ADP-forming) [EC:6.2.1.13]; acs, acetyl-CoA synthase [EC:2.3.1.169]; *atoAD*, acetate CoA/acetoacetate CoA-transferase [EC:2.8.3.8 2.8.3.9]; EC:6.2.1.2, medium-chain acyl-CoA synthetase [EC:6.2.1.2]; *fadD*, long-chain acyl-CoA synthetase [EC:6.2.1.3]; *pccA*, propionyl-CoA carboxylase alpha chain [EC:6.4.1.3]; *epi*, methylmalonyl-CoA/ethylmalonyl-CoA epimerase [EC:5.1.99.1]; *mut*, methylmalonyl-CoA mutase [EC:5.4.99.2]; *mcl*, malyl-CoA/(S)-citramalyl-CoA lyase [EC:4.1.3.24 4.1.3.25]; *mch*, 2-methylfumaryl-CoA hydratase [EC:4.2.1.148]; *mct*, 2-methylfumaryl-CoA isomerase [EC:5.4.1.3]; *meh*, 3-methylfumaryl-CoA hydratase [EC:4.2.1.153]; *smtAB*, succinyl-CoA:(S)-malate CoA-transferase subunit A [EC:2.8.3.22]; *prpB*, methylisocitrate lyase [EC:4.1.3.30]; *prpC*, 2-methylcitrate synthase [EC:2.3.3.5]; *prpD*, 2-methylcitrate dehydratase [EC:4.2.1.79]; *bcd*, butyryl-CoA dehydrogenase [EC:1.3.8.1]; *acd*, acyl-CoA dehydrogenase [EC:1.3.8.7]; *paaF*, enoyl-CoA hydratase [EC:4.2.1.17]; *crt*, enoyl-CoA hydratase [EC:4.2.1.17]; *paaH*, 3-hydroxybutyryl-CoA dehydrogenase [EC:1.1.1.157]; *phbB*, acetoacetyl-CoA reductase [EC:1.1.1.36]; *atoB*, acetyl-CoA C-acetyltransferase [EC:2.3.1.9]; *fadJ*, 3-hydroxyacyl-CoA dehydrogenase / enoyl-CoA hydratase / 3-hydroxybutyryl-CoA epimerase [EC:1.1.1.35 4.2.1.17 5.1.2.3]; *fadA*, acetyl-CoA acyltransferase [EC:2.3.1.16]; *dehH*, 2-haloacid dehalogenase [EC:3.8.1.2]; EC:3.8.1.3, haloacetate dehalogenase [EC:3.8.1.3]; *glcDEF*, glycolate oxidase [EC:1.1.3.15]; EC:1.1.3.15, (S)-2-hydroxy-acid oxidase [EC:1.1.3.15].

Alcohol and aldehyde dehydrogenases sequentially oxidize the resulting alcohols to their corresponding fatty acids or fatty acyl-CoA. Binatota genomes encode a plethora of alcohol and aldehyde dehydrogenases. These include the wide substrate range alcohol (EC:1.1.1.1), and aldehyde (EC:1.2.1.3) dehydrogenases encoded by the majority of Binatota genomes, as well as bifunctional alcohol/aldehyde dehydrogenase (EC:1.2.1.10 /1.1.1.1) encoded by a few Binatota genomes (7 genomes), and some highly specific enzymes, e.g. the short-chain isopropanol dehydrogenase (EC:1.1.1.80) for converting isopropanol and other secondary alcohols to the corresponding ketone (20 genomes), and acetone monooxygenase (*acmA*, EC:1.14.13.226) and methyl acetate hydrolase (*acmB*, EC:3.1.1.114) that will sequentially oxidize acetone to methanol and acetate (6 genomes) (Figure 4a, Extended data 1).

A Complete fatty acid degradation machinery that enables all orders of the Binatota to degrade short-, medium-, and long-chain fatty acids to acetyl CoA and propionyl-CoA were identified (Figure 4b, Extended data 1). Acetyl-CoA produced from the beta-oxidation pathway could be assimilated via the ethylmalonyl CoA pathway (EMCP) or the glyoxylate shunt as discussed above. Further, two pathways for propionyl-CoA assimilation, generated from the degradation of odd chain fatty acids, were identified (Figure 4c). Orders Bin18, Binatales, UBA1149, UBA12105, and UTPR01 all encode enzymes for the methylmalonyl CoA (MMCoA) pathway that carboxylates propionyl CoA to succinyl-CoA (TCA cycle intermediate) via a methylmalonyl-CoA intermediate. On the other hand, the majority of order UBA9968 genomes encode enzymes of the 2-methylcitrate cycle for propionyl-CoA degradation (*prpBCD*) where propionate is degraded to pyruvate and succinate via a 2-methylcitrate intermediate (Figures 4b-c, Extended data 1).

### Electron transport chain

All Binatota genomes encode an aerobic respiratory chain comprising complexes I, II, and IV, as well as an F-type H^+^-translocating ATP synthase (Figures 5a-b, Extended data 1). Interestingly, genes encoding complex III (cytochrome bc1 complex) were sparse in Binatota genomes with some orders lacking genes encoding all subunits (e.g. HRBin30) and others only encoding the Fe-S (ISP) and the cytochrome b (*cytB*) but not the cytochrome c1 (*cyt1*) subunit (e.g. Binatales, UBA1149). Instead, genes encoding an Alternate Complex III (ACIII, encoded by *actABCDEFG*) were identified in 76 genomes, with 12 genomes encoding both complete complexes (in orders Bin 18, UBA9968, and UTPR01). Complex III and ACIII transfer electrons from reduced quinones (all genomes encode the capability of menaquinone biosynthesis) to cytochrome c which, in turn, reduces cytochrome c oxidase (complex IV). Homologues of the electron transfer proteins belonging to cytochrome c families were rare in Binatota genomes, especially those encoding ACIII (Figure 5a, Extended data 1). However, the recent structure of ACIII from *Flavobacterium johnsoniae* ^42^ in a supercomplex with cytochrome c oxidase aa3 suggests that electrons could potentially flow from ACIII to complex IV without the need for cytochrome c, which might explain the paucity of cytochrome c homologues in ACIII-harboring genomes.

**Figure 5.**
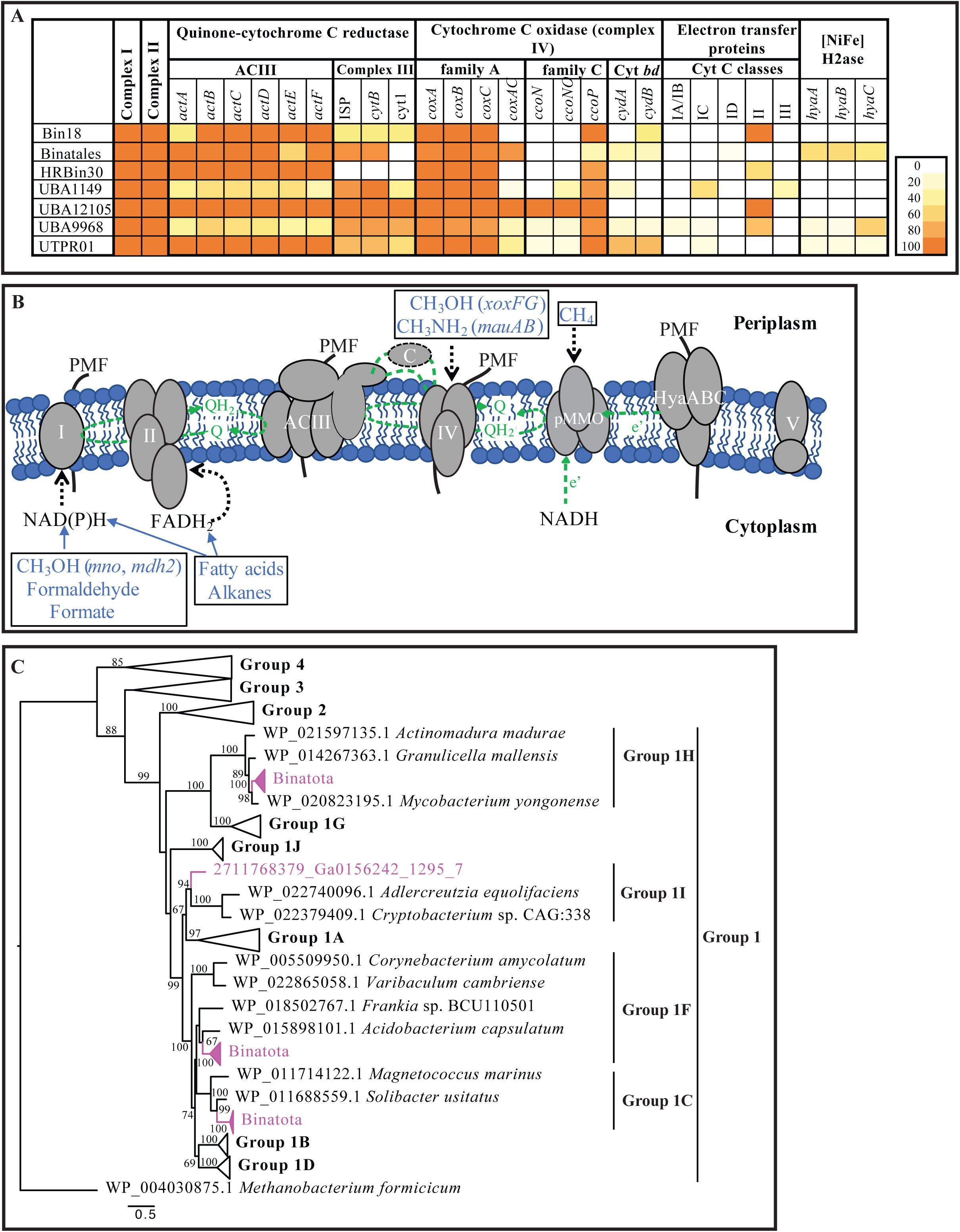
Electron transport chain in the Binatota. (A) Heatmap of the distribution of electron transport chain components in the Binatota genomes and electrons entry points from various substrates. The heatmap colors (as explained in the key) correspond to the percentage of genomes in each order encoding a homologue of the gene in the column header. All subunits of complexes I (NADH-quinone oxidoreductase [EC:7.1.1.2]), and II (succinate dehydrogenase /fumarate reductase [EC:1.3.5.1 1.3.5.4]) were encoded in all genomes but are shown here as single components for ease of visualization. Genes encoding quinone-cytochrome C reductase activities belonged to either complex III (cytochrome bc1; ISP/*cytb*/cyt1) and/or alternate cytochrome III (ACIII; *actABCDEF*), while genes encoding cytochrome c oxidase activities (complex IV) belonged to different families including family A (cytochrome c oxidase aa3; *coxABC*), family C (cytochrome c oxidase cbb3; *ccoNOP*), and/or cytochrome *bd* (*cydAB*). Possible electron transfer proteins between complex III (or alternate complex III) and complex IV belonging to different cytochrome c families are shown. Also shown in (A) is the distribution of the three subunits of the type I respiratory O_2_-tolerant H_2_-uptake [NiFe] hydrogenase (*hyaABC*) in Binatota genomes. (B) A cartoon depicting all electron transfer complexes (I, II, ACIII, IV) embedded in the inner membrane, along with the particulate methane monooxygenase (pMMO), and the H_2_-uptake [NiFe] hydrogenase (HyaABC). All genomes also encoded an F-type ATP synthase complex (V). Substrates potentially supporting growth are shown in blue with predicted entry points to the ETC shown as dotted black arrows. Sites of proton extrusion to the periplasm and PMF creation are shown as solid black lines, while sites of electron (e’) transfer are shown as dotted green lines. Three possible physiological reductants are shown for pMMO (as dotted green arrows); the quinone pool coupled to ACIII, NADH, and/or some of the reduced quinones generated through H_2_ oxidation by HyaABC. (C) Maximum likelihood phylogenetic tree showing the classification of the *hyaA* genes encoded by the Binatota genomes (magenta) in relation to other [Ni-Fe] hydrogenases. The [Fe-Fe] hydrogenase of *Methanobacterium formicum* was used as the outgroup. Bootstrap support (from 100 bootstraps) is shown for branches with >50% support.

Based on the predicted ETC structure, the flow of electrons under different growth conditions in the Binatota could be envisaged (Figure 5b). When growing on methane, pMMO would be coupled to the electron transport chain at complex III level via the quinone pool, where reduced quinones would act as physiological reductant of the enzyme ^43^ (Figure 5b). pMMO was also previously reported to receive electrons donated by NADH ^44^. During methanol oxidation by periplasmic enzymes (e.g. *xoxF*-type methanol dehydrogenases), and methylamine oxidation by the periplasmic methylamine dehydrogenase (*mauAB*) electrons would be shuttled via their respective C-type cytochrome (*xoxG*, and *mauC*, respectively) to complex IV. In the cytosol, methanol oxidation via the *mno/mdo*-type or the *mdh2*-type methanol dehydrogenases, as well as formaldehyde and formate oxidation via the action of cytoplasmic formaldehyde and formate dehydrogenases would contribute NADH to the aerobic respiratory chain through complex I. Similarly, when growing heterotrophically on alkanes and/or fatty acids, reducing equivalents in the form of NAD(P)H, and FADH_2_ serve as electron donors for aerobic respiration through complex I, and II, respectively (Figure 5b).

Binatota genomes also encode respiratory O_2_-tolerant H_2_-uptake [NiFe] hydrogenases, belonging to groups 1c (6 sequences), 1f (22 sequences), 1i (1 sequence), and 1h (4 sequences) (Figure 5c). In *E. coli*, these membrane-bound periplasmically oriented hydrogenases transfer electrons (through their cytochrome b subunit) from molecular hydrogen to the quinone pool. Cytochrome *bd* oxidase (complex IV) then completes this short respiratory electron transport chain between H_2_ and O_2_ ^45^. In *E. coli*, the enzyme functions under anaerobic conditions ^46^, and may function as an O_2_-protecting mechanism ^47^. Further, simultaneous oxidation of hydrogen (via type I respiratory O_2_-tolerant hydrogenases) and methane (via pMMO) has been shown to occur in methanotrophic Verrucomicrobia to maximize proton-motive force generation and subsequent ATP production ^48^. As well, some of the reduced quinones generated through H_2_ oxidation are thought to provide reducing power for catalysis by pMMO ^48^ (Figure 5b).

### Pigment production genes in the Binatota

*Carotenoids.* Analysis of the Binatota genomes demonstrated a wide range of hydrocarbon (carotenes) and oxygenated (xanthophyll) carotenoid biosynthesis capabilities. Carotenoids biosynthetic machinery in the Binatota included *crtB* for 15-cis-phyotene synthesis from geranylgeranyl-PP; *crtI*, *crtP*, *crtQ*, and *crtH* for neurosporene and all-*trans* lycopene formation from 15-cis-phytone; *crtY* or *crtL* for gamma-and beta-carotene formation from all-*trans* lycopene; and a wide range of genes encoding enzymes for the conversion of neurosporene to spheroidene and 7,8-dihydro β-carotene, as well as the conversion of all-trans lycopene to spirilloxanthin, gamma-carotene to hydroxy-chlorobactene glucoside ester and hydroxy-Ɣ-carotene glucoside ester, and beta carotene to isorenieratene and zeaxanthins (Figures 6a-b, Extended data 1). Gene distribution pattern (Figure 6a, Extended data 1) predicts that all Binatota orders are capable of neurosporene and all-trans lycopene biosynthesis, and all but the order HRBin30 are capable of isorenieratene, zeaxanthin, β-carotene and dihydro β-carotene biosynthesis, and with specialization of order UTPR01 in spirilloxanthin, spheroidene, hydroxy-chlorobactene, and hydroxy Ɣ-carotene biosynthesis.

**Figure 6.**
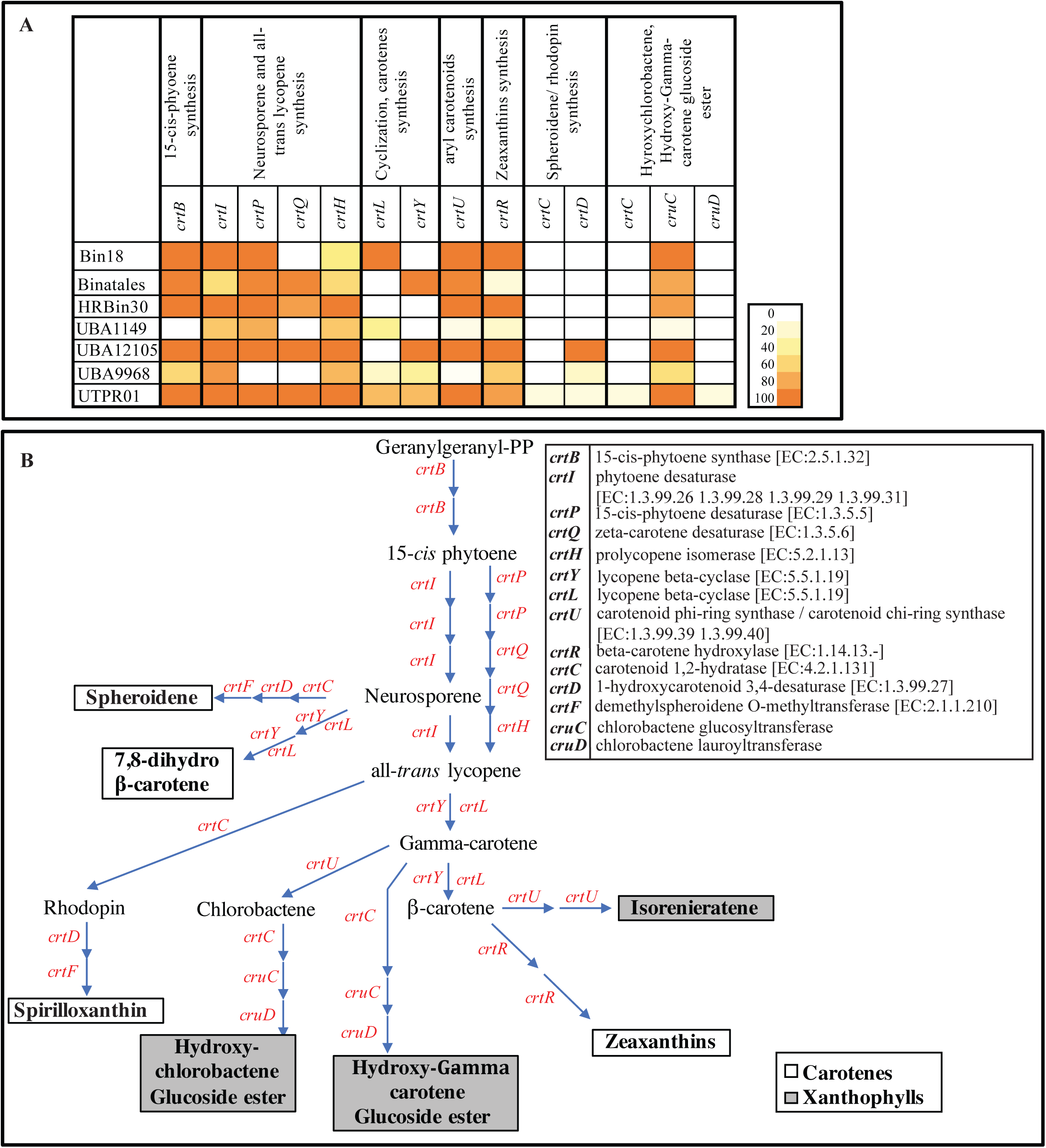
Carotenoids biosynthesis capabilities in Binatota genomes. (A) Distribution of carotenoid biosynthesis genes in the Binatota genomes. The heatmap colors (as explained in the key) correspond to the percentage of genomes in each order encoding a homologue of the gene in the column header. (B) Carotenoid biosynthesis scheme in Binatota based on the identified genes. Genes encoding enzymes catalyzing each step are shown in red and their descriptions with EC numbers are shown to the right. Binatota genomes encode the capability to biosynthesize both exclusively hydrocarbon carotenes (white boxes), or the oxygenated xanthophylls (grey boxes).

### Bacteriochlorophylls

Surprisingly, homologues of multiple genes involved in bacteriochlorophyll biosynthesis were ubiquitous in Binatota genomes (Figure 7a-c). Bacteriochlorophyll biosynthesis starts with the formation of chlorophyllide *a* from protoporphyrin IX (Figure 7b). Within this pathway, genes encoding the first *bchI* (Mg-chelatase [EC:6.6.1.1]), third *bchE* (magnesium-protoporphyrin IX monomethyl ester cyclase [EC:1.21.98.3]), and fourth *bchLNB* (3,8-divinyl protochlorophyllide reductase [EC:1.3.7.7]) steps were identified in the Binatota genomes (Figures 7a, 7b, Extended data 1). However, homologues of genes encoding the second *bchM* (magnesium-protoporphyrin O-methyltransferase [EC:2.1.1.11]), and the fifth (*bciA* or *bicB* (3,8-divinyl protochlorophyllide *a* 8-vinyl-reductase), or *bchXYZ* (chlorophyllide *a* reductase, EC 1.3.7.15])) steps were absent (Figure 7a-b). A similar patchy distribution was observed in the pathway for bacteriochlorophyll *a* (Bchl *a*) formation from chlorophyllide *a* (Figure 7b), where genes encoding *bchXYZ* (chlorophyllide *a* reductase [EC 1.3.7.15]) and *bchF* (chlorophyllide *a* 3^1^-hydratase [EC 4.2.1.165]) were not identified, while genes encoding *bchC* (bacteriochlorophyllide *a* dehydrogenase [EC 1.1.1.396]), *bchG* (bacteriochlorophyll a synthase [EC:2.5.1.133]), and *bchP* (geranylgeranyl-bacteriochlorophyllide *a* reductase [EC 1.3.1.111)) were present in most genomes (Figure 7a, Extended data 1). Finally, within the pathway for bacteriochlorophylls *c* (Bchl *c*) and *d* (Bchl *d*) formation from chlorophyllide *a* (Figure 7b), genes for *bciC* (chlorophyllide a hydrolase [EC:3.1.1.100]), and *bchF* (chlorophyllide *a* 3^1^-hydratase [EC:4.2.1.165]) or *bchV* (3-vinyl bacteriochlorophyllide hydratase [EC:4.2.1.169] were not identified, while genes for *bchR* (bacteriochlorophyllide d C-12(1)-methyltransferase [EC:2.1.1.331]), *bchQ* (bacteriochlorophyllide d C-8(2)-methyltransferase [EC:2.1.1.332]), *bchU* (bacteriochlorophyllide d C-20 methyltransferase [EC:2.1.1.333]), and *bchK* (bacteriochlorophyll c synthase [EC:2.5.1.-]) were identified (Figure 7b, Extended data 1).

**Figure 7.**
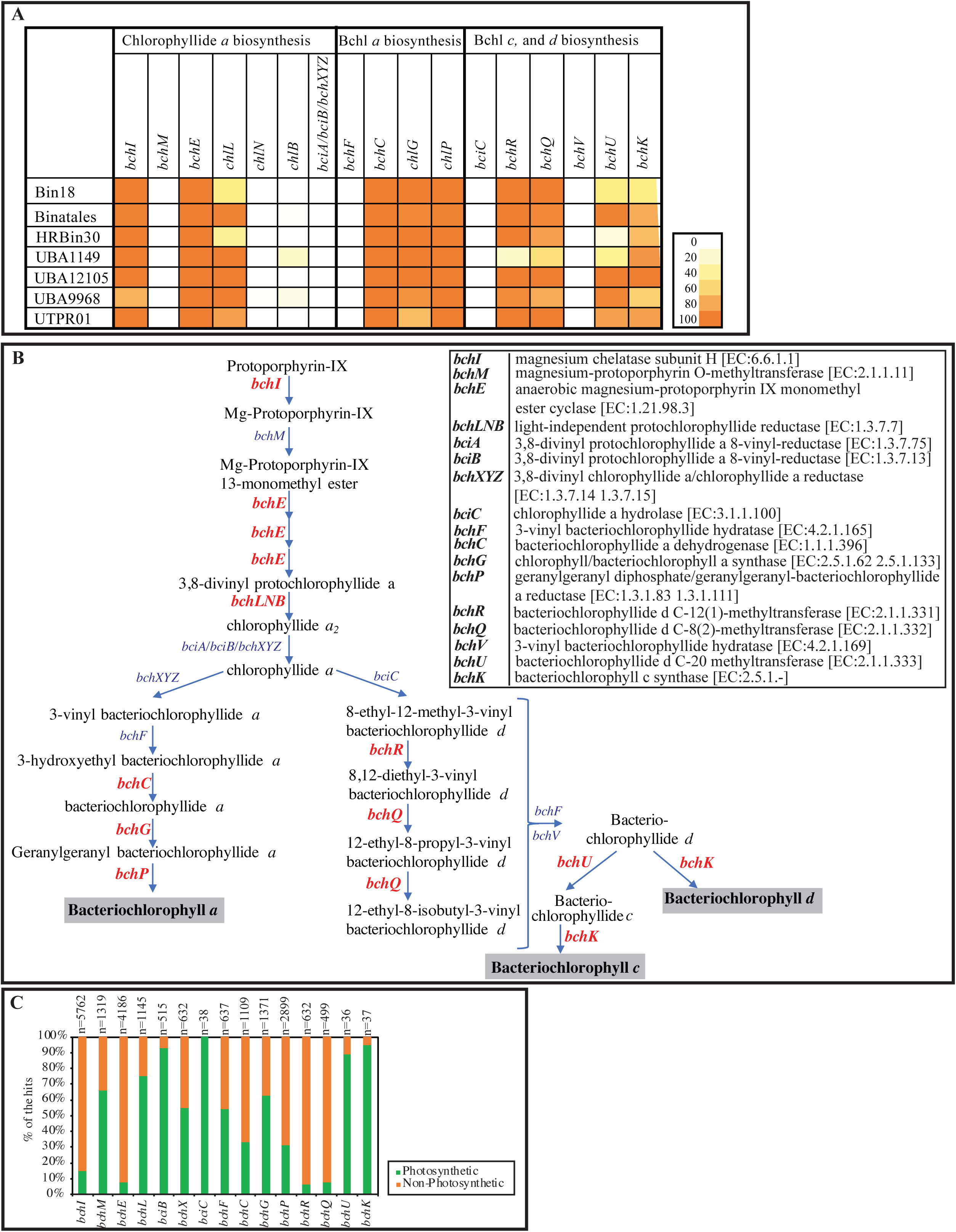
Bacteriochlorophylls biosynthesis genes encountered in Binatota genomes studied suggesting an incomplete pathway for bacteriochlorophyll *a*, *c*, and/or *d* biosynthesis. (A) Distribution of chlorophyll biosynthesis genes in Binatota genomes. The heatmap colors (as explained in the key) correspond to the percentage of genomes in each order encoding a homologue of the gene in the column header. (B) Bacteriochlorophylls biosynthesis pathway. Genes identified in at least one Binatota genome are shown in red boldface text, while these with no homologues in the Binatota genomes are shown in blue text. Gene descriptions with EC numbers are shown to the right of the figure. (C) Distribution patterns of bacteriochlorophyll biosynthesis genes. The search was conducted in the functionally annotated bacterial tree of life AnnoTree ^96^ using single KEGG orthologies implicated in chlorophyll biosynthesis. Gene names are shown on the X-axis, total number of hits are shown above the bars for each gene, and the percentage of hits in genomes from photosynthetic (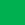) versus non-photosynthetic (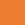) genera are in the stacked bars.

### Ecological distribution of the Binatota

A total of 1,889 (GenBank nt) and 1,213 (IMG/M) 16S rRNA genes affiliated with the Binatota orders were identified (Extended data 2 and 3, Figures 8, S1a). Analyzing their environmental distribution showed preference of Binatota to terrestrial soil habitats (39.5-83.0% of GenBank, 31.7-91.6% of IMG/M 16S rRNA gene sequences in various orders), as well as plant-associated (particularly rhizosphere) environments; although this could partly be attributed to sampling bias of these globally distributed and immensely important ecosystems (Figure 8a). On the other hand, a paucity of Binatota-affiliated sequences was observed in marine settings, with sequences absent or minimally present for Binatales, HRBin30, UBA9968, and UTPRO1 datasets (Figure 8a). The majority of sequences from marine origin were sediment-associated, being encountered in hydrothermal vents, deep marine sediments, and coastal sediments, with only the Bin18 sequences sampled from IMG/M showing representation in the vast, relatively well-sampled pelagic waters (Figure 8d).

**Figure 8.**
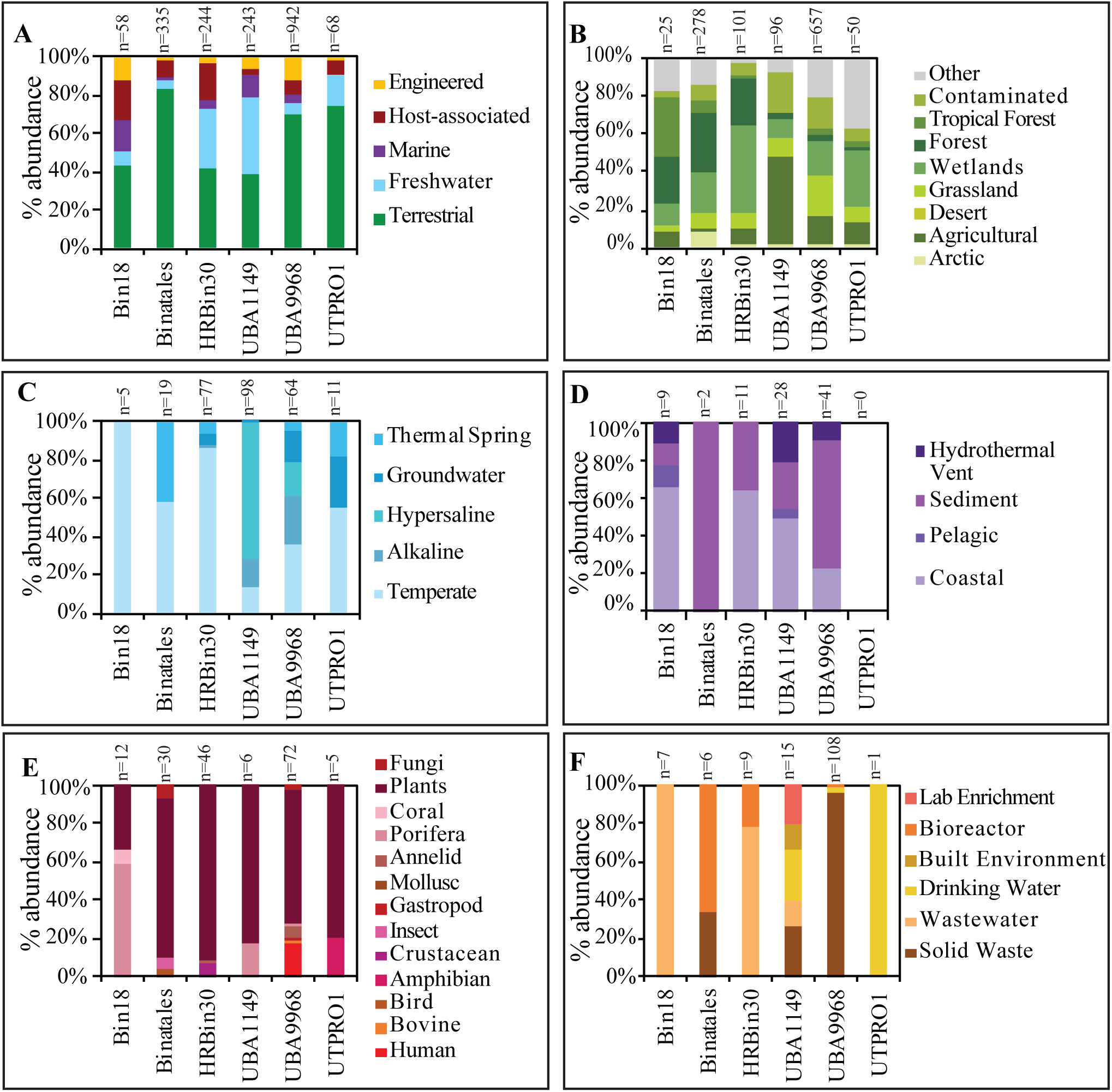
Ecological distribution of Binatota-affiliated 16S rRNA sequences in GenBank nt database. Binatota orders are shown on the X-axis, while percentage abundance in different environments (classified based on the GOLD ecosystem classification scheme) are shown on the Y-axis (A). Further sub-classifications for each environment are shown for (B) terrestrial, (C) freshwater, (D) marine, (E) host-associated, and (F) engineered environments. The total number of hit sequences for each order are shown above the bar graphs. Details including GenBank accession number of hit sequences are shown in Extended Data 2. Order UBA12015 genome assembly did not contain a16S rRNA gene, and so this order is not included in the analysis.

In addition to phylum-wide patterns, order-specific environmental preferences were also observed. For example, in order Bin18, one of the two available genomes originated from the Mediterranean sponge *Aplysina aerophoba*. Analysis of the 16S rRNA dataset suggests a notable association between Bin18 and sponges, with a relatively high host-associated sequences (Figure 8a), the majority of which (58.3% NCBI-nt, 25.0% IMG/M) were recovered from the Porifera microbiome (Figures 8e, S1f). Bin18-affiliated 16S rRNA gene sequences were identified in a wide range of sponges from ten genera and five global habitat ranges (the Mediterranean genera *Ircinia*, *Petrosia*, *Chondrosia*, and *Aplysina*, the Caribbean genera *Agelas*, *Xestospongia*, and *Aaptos*, the Indo-West Pacific genus *Theonella*, the Pacific Dysideidae family, and the Great Barrier Reef genus *Rhopaloeides*), suggesting its widespread distribution beyond a single sponge species. The absolute majority of order Binatales sequences (83.0% NCBI-nt, 91.6% IMG/M) were of a terrestrial origin (Figures 8a, S1c), in addition to multiple rhizosphere-associated samples (7.5% NCBI-nt and 2.8% IMG/M, respectively) (Figure 8a, S1f). Notably, a relatively large proportion of Binatales soil sequences originated either from wetlands (peats, bogs) or forest soils (Figures 8b, S1c), strongly suggesting the preference of the order Binatales to acidic and organic/methane-rich terrestrial habitats. This corresponds with the fact that 42 out of 48 Binatales genomes were recovered from soil, 38 of which were from acidic wetland or forest soils (Figure 1, Table S1). Genomes of UBA9968 were recovered from a wide range of terrestrial and non-marine aquatic environments, and the observed 16S rRNA gene distribution enforces their ubiquity in all but marine habitats (Figures 8a, S1b-g). Finally, while genomes from orders HRBin30, UBA1149 and UTPRO1 were recovered from limited environmental settings (thermal springs for HRBin30, gaseous hydrocarbon impacted habitats, e.g. marine hydrothermal vents and gas-saturated Lake Kivu for UBA1149, and soil and hydrothermal environments for UTPRO1) (Figure 1, Table S1), 16S rRNA gene analysis suggested their presence in a wide range of environments from each macro-scale environment classification (Figures 8a, S1b-g).

## Discussion

### Expanding the world of methylotrophy

The current study expands the list of lineages potentially capable of methylotrophy. An extensive repertoire of genes and pathways mediating the oxidation of multiple C1 compounds to formaldehyde (Figure 2, 9), formaldehyde oxidation to CO_2_ (Figure 3a), as well as formaldehyde assimilation pathways (Figure 3c) were identified, indicating that such capacity is a defining metabolic trait in the Binatota. A certain degree of order-level substrate preference was observed, with potential utilization of methanol in all orders except HRBin30, methylamine in all orders except UBA9968, S-containing C1 compound in Bin18, Binatales, and UBA9968, halogenated methane in Bin18, and possible methane utilization (methanotrophy) in Bin18 and Binatales (Figure 2a).

**Figure 9.**
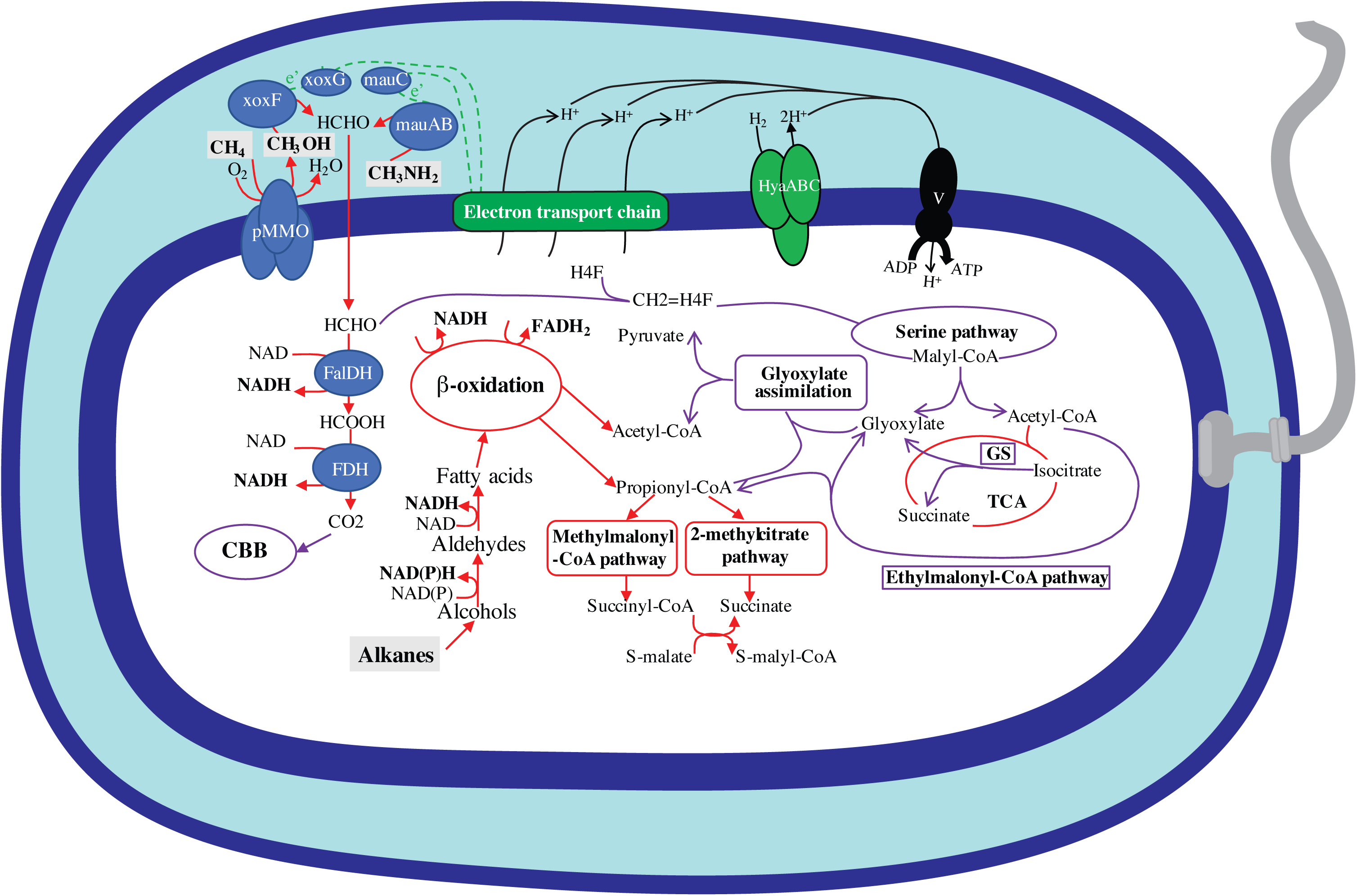
Cartoon depicting different metabolic capabilities encoded in the Binatota genomes. Enzymes for C1 metabolism are shown in blue and include the periplasmic particulate methane monooxygenase (pMMO), methanol dehydrogenase (*xoxFG*), and methylamine dehydrogenase (*mauABC*), as well as the cytoplasmic formaldehyde dehydrogenase (FalDH), and formate dehydrogenase (FDH). Electron transport chain is shown as a green rectangle. Electron transfer from periplasmic enzymes to the ETC is shown as dotted green lines (details of the ETC are shown in Figure 5b). The sites of proton extrusion to the periplasm are shown as black arrows, as is the F-type ATP synthase. Carbon dissimilation routes are shown as red arrows, while assimilatory routes are shown as purple arrows. Details of the assimilatory pathways are shown in Figures 2 and 3. Reducing equivalents potentially fueling the ETC (NAD(P)H, and FADH_2_) are shown in boldface. All substrates predicted to support growth are shown in boldface within grey boxes. A flagellum is also depicted, the biosynthetic genes of which were identified in genomes belonging to all orders except Bin18, HRBin30, and UBA1149. The cell is also depicted as rod-shaped based on the identification of the rod shape determining protein *rodA* in all genomes, and the rod-shape determining proteins *mreB* and *mreC* in genomes from all orders except UBA1149. Abbreviations: CBB, Calvin Bensen Basham cycle; Fal-DH, NAD-linked glutathione-independent formaldehyde dehydrogenase, *fdhA*; FDH, NAD-dependent formate dehydrogenase [EC: 1.17.1.9); Fum, fumarate; GS, glyoxylate shunt; H_4_F, tetrahydrofolate; HyaABC, type I respiratory O_2_-tolerant H_2_-uptake [NiFe] hydrogenase; *mauABC*, methylamine dehydrogenase; *pmoABC*, particulate methane monooxygenase; *xoxFG*, xoxF-type methanol dehydrogenase; succ, succinate; TCA, tricarboxylic acid cycle; V, F-type ATP synthase [EC:7.1.2.2 7.2.2.1].

Aerobic methylotrophy has been documented in members of the alpha, beta, and gamma Proteobacteria ^49^, Bacteroidetes ^50^, Actinobacteria (e.g. genera *Arthrobacter* and *Mycobacterium*), Firmicutes (e.g. *Bacillus methanolicus*) ^51^, Verrucomicrobia ^52^, and *Candidatus* Methylomirabilis (NC10) ^53^. Further, studies employing genome-resolved metagenomics identified some signatures of methylotrophy, e.g. methanol oxidation ^10, 54^, formaldehyde oxidation/ assimilation ^55^, and methylamine oxidation ^10^, in the Gemmatimonadetes, Rokubacteria, Chloroflexi, Actinobacteria, Acidobacteria, and Lambdaproteobacteria. The possible contribution of Binatota to methane oxidation (methanotrophy) is especially notable, given the global magnitude of methane emissions ^56^, and the relatively narrower range of organisms (Proteobacteria, Verrucomicrobia, and *Candidatus* Methylomirabilota (NC10)) ^57^ capable of this special type of methylotrophy. As described above, indirect evidence exists for the involvement of Binatota harboring TUSC-type pMMO sequences in methane oxidation, while it is currently uncertain whether Binatota harboring SAR324/Actinobacteria-type pMMO sequences are involved in oxidation of methane, gaseous alkanes, or both. pMMO of methanotrophs is also capable of oxidizing ammonia to hydroxylamine, which necessitates methanotrophs to employ hydroxylamine detoxification mechanisms ^58^. All eleven Binatota genomes encoding pMMO also encoded at least one homologue of *nir*, *nor*, and/or *nos* genes that could potentially convert harmful N-oxide byproducts to dinitrogen.

As previously noted ^22^, methylotrophy requires the possession of three metabolic modules: C1 oxidation to formaldehyde, formaldehyde oxidation to CO_2_, and formaldehyde assimilation. Within the world of methylotrophs, a wide array of functionally redundant enzymes/pathways has been characterized that mediates various reactions/ transformations in such modules. In addition, multiple combinations of different modules have been observed in methylotrophs, with significant variations existing even in phylogenetically related organisms. Our analysis demonstrates that such metabolic versatility indeed occurs within Binatota’s methylotrophic modules. While few phylum-wide characteristics emerged, e.g. utilization of serine pathway for formaldehyde assimilation, absence of H_4_MPT-linked formaldehyde oxidation, and potential utilization of PEP carboxykinase (*pckA*) rather than PEP carboxylase (*ppc*) for CO_2_ entry to the serine cycle, multiple order-specific differences were observed, e.g. XoxF-type methanol dehydrogenase encoded by Bin18 and Binatales genomes, MDH2-type methanol dehydrogenase encoded by UBA1149 genomes, absence of methanol dehydrogenase homologues in HRBin30 genomes, absence of methylamine oxidation in order UBA9968, and potential utilization of the ethylmalonyl-CoA pathway for glyoxylate regeneration by the majority of the orders versus the glyoxylate shunt by UBA9968.

### Alkane degradation in the Binatota

A second defining feature of the phylum Binatota, besides methylotrophy, is the widespread capacity for aerobic alkane degradation, as evident by the extensive arsenal of genes mediating aerobic degradation of short-(pMMO, propane monooxygenase), medium-(*alkB*, cyp153), and long-chain alkanes (*ladA*) identified (Figure 4a), in addition to complete pathways for odd-and even-numbered fatty acids oxidation (Figure 4b). Hydrocarbons, including alkanes, have been an integral part of the earth biosphere for eons, and a fraction of microorganisms has evolved specific mechanisms (O_2_-dependent hydroxylases and monooxygenases, anaerobic addition of fumarate) for their activation and conversion to central metabolites ^59^. Aerobic alkane degradation capacity has so far been encountered in the Actinobacteria, Proteobacteria, Firmicutes, Bacteroidetes, as well as in a few Cyanobacteria ^59^. As such, this study adds to the expanding list of phyla capable of aerobic alkane degradation.

### Metabolic traits explaining niche preferences in the Binatota

Analysis of 16S rRNA gene datasets indicated that the Binatota display phylum-wide (preference to terrestrial habitats and methane/hydrocarbon-impacted habitats, and rarity in pelagic marine environments), as well as order-specific (Bin18 in sponges, HRBin30 and UBA1149 in geothermal settings, Binatales in peats, bogs, and forest soils) habitat preferences (Figures 8, S1). Such distribution patterns could best be understood in light of the phylum’s predicted metabolic capabilities. Soils represent an important source of methane, generated through microoxic and anoxic niches within soil’s complex architecture ^60^. Methane emission from soil is especially prevalent in peatlands, bogs, and wetlands, where incomplete aeration and net carbon deposition occurs. Indeed, anaerobic ^61^, fluctuating ^62^, and even oxic ^63^ wetlands represent one of the largest sources of methane emissions to the atmosphere. As well, terrestrial ecosystems represent a major source of global methanol emissions ^64^, with its release mostly mediated by demethylation reactions associated with pectin and other plant polysaccharides degradation. C1-metabolizing microorganisms significantly mitigate methane and methanol release to the atmosphere from terrestrial ecosystems ^65^, and we posit that members of the Binatota identified in soils, rhizosphere, and wetlands contribute to such process. The special preference of order Binatales to acidic peats, bogs, forests, and wetlands could reflect a moderate acidophilic specialization for this order and suggest their contribution to the process in these habitats.

Within the phylum Binatota, it appears that orders HRBin30 and UBA1149 are abundant in thermal vents, thermal springs, and thermal soils, suggesting a specialization to high temperature habitats (Figure 8). Binatota’s presence in such habitats could be attributed to high concentrations of alkanes typically encountered in such habitats. Hydrothermal vents display steep gradients of oxygen in their vicinity, emission of high levels of methane and other gaseous alkanes, as well as thermogenic generation of medium-and long-chain alkanes ^66^. Indeed, the presence and activity of aerobic hydrocarbon degraders in the vicinity of hydrothermal vents have been well established ^30, 31, 67^.

The recovery of Binatota genomes from certain lakes could be a reflection of the high gaseous load in such lakes. Multiple genomes and a large number of Binatota-affiliated 16S rRNA sequences were binned/identified from Lake Kivu, a meromictic lake characterized by unusually high concentrations of methane ^68^. Biotically, methane evolving from Lake Kivu is primarily oxidized by aerobic methanotrophs in surface waters ^68, 69, 70^, and members of the Binatota could contribute to this process. Binatota genomes were also recovered from Lake Washington sediments, a location that has long served as a model for studying methylotrophy ^71, 72^. Steep counter gradients of methane and oxygen occurring in the Lake’s sediments enable aerobic methanotrophy to play a major role in controlling methane flux through the water column ^73, 74, 75, 76^.

Finally, the occurrence and apparent wide distribution of members of the Binatota in sponges, particularly by the order Bin18, is notable, and could be attributed to the recognized nutritional-based symbiosis between sponges and hydrocarbon-degraders ^77, 78^, including methanotrophs ^79^. This is especially true in deep-water sponges, where low levels of planktonic biomass restrict the amount of food readily acquired via filter feeding and hence biomass acquisition via methane and alkane oxidation is especially valuable.

### Carotenoid pigmentation: occurrence and significance

The third defining feature of the Binatota, in addition to aerobic methylotrophy and alkane degradation, is the predicted capacity for carotenoid production. In photosynthetic organisms, carotenoids increase the efficiency of photosynthesis by absorbing in the blue-green region then transferring the absorbed energy to the light-harvesting pigments ^80^. Carotenoid production also occurs in a wide range of non-photosynthetic bacteria belonging to the Alpha-, Beta, and Gamma-Proteobacteria (including methano-and methylotrophs ^81^), Bacteroidetes ^82^, Deinococcus ^83^, Thermus ^84^, Delta-Proteobacteria ^85^, Firmicutes ^86^, Actinobacteria ^87^, Planctomycetes ^88^, and Archaea, e.g. Halobacteriaceae ^89^, and *Sulfolobus* ^90^. Here, carotenoids could serve as antioxidants ^91^, and aid in radiation, UV, and desiccation resistance ^92, 93^. The link between carotenoid pigmentation and methylo/methanotrophy has long been observed ^94^, with the majority of known model Alpha- and Gamma-Proteobacteria methano- and methylotrophs being carotenoid producers, although several Gram-positive methylotrophs (*Mycobacterium*, *Arthrobacter*, and *Bacillus*) are not pigmented. Indeed, root-associated facultative methylotrophs of the genus *Methylobacterium* have traditionally been referred to as “pink pigmented facultative methylotrophs” and are seen as integral part of root ecosystems ^95^. The exact reason for this correlation is currently unclear and could be related to the soil environment where they are prevalent, where periodic dryness and desiccation could occur, or to the continuous exposure of these aerobes in some habitats to light (e.g. in shallow sediments), necessitating protection from UV exposure.

### Chlorophyll biosynthesis genes in the Binatota

Perhaps the most intriguing finding in this study is the identification of the majority of genes required for the biosynthesis of bacteriochlorophylls from protoporphyrin-IX (six out of ten genes for bacteriochlorophyll *a* and seven out of eleven genes for bacteriochlorophyll *c* and *d*). While such pattern is tempting to propose phototrophic capacities in the Binatota based on the common practice of using a certain percentage completion threshold to denote pathway occurrence in some studies (e.g. ^2^), the consistent absence of critical genes (*bchM* methyltransferase, *bciA*/*bciB*/*bchXYZ* reductases, *bciC* hydrolase, and *bchF*/*V* hydratases), coupled with our inability to detect reaction center-encoding genes, prevents such a proclamation. Identification of a single or few gene shrapnel from the chlorophyll biosynthesis pathway in microbial genomes is not unique. Indeed, searching the functionally annotated bacterial tree of life AnnoTree ^96^ using single KEGG orthologies implicated in chlorophyll biosynthesis identifies multiple (in some cases thousands) hits in genomes from non-photosynthetic organisms (Figure 7c). This is consistent with the identification of a *bchG* gene in a Bathyarchaeota fosmid clone ^97^, and, more recently, a few bacteriochlorophyll synthesis genes in an Asgard genome ^98^. However, it should be noted that the high proportion of genes in the bacteriochlorophyll biosynthetic pathway identified in the Binatota genomes has never previously been encountered in non-photosynthetic microbial genomes. Indeed, a search in AnnoTree for the combined occurrence of all seven bacteriochlorophyll synthesis genes identified in Binatota genomes yielded only photosynthetic organisms.

Accordingly, we put forward three scenarios to explain the proposed relationship between Binatota and phototrophy: The most plausible scenario, in our opinion, is that members of the Binatota are pigmented non-photosynthetic organisms capable of carotenoid production, but incapable of chlorophyll production and lack a photosynthetic reaction center. Under this scenario, the incomplete pathway for bacteriochlorophyll biosynthesis represents a pattern of gene loss from a chlorophyll-producing ancestor. The assumption that a lineage has lost the immensely beneficial capacity to harvest energy from light might appear counterintuitive, even implausible. However, this could be understood in the context of the proposed role of chlorophyll during the early evolution of photosynthesis. In a thought-provoking review, Martin et al. ^99^ argue that the evolution of chlorophyll-based biosynthesis occurred against a backdrop of chemolithotrophy in hydrothermal vents, with hydrogen produced abundantly by serpentinization as the main source of energy, and CO_2_ fixation via the acetyl CoA pathway as the main source of carbon. The acetyl-CoA pathway requires electron transfer to an acceptor, ferrodoxin, with an extremely negative midpoint potential, which could only be achieved via electron bifurcation reactions. Within such chemolithotrophic, dim-lit hydrogen-dominated realm, the main benefit of chlorophyll-based anoxygenic photosynthesis would be harvesting the relatively limited amount of thermal light emitted from hydrothermal vents ^100^ to allow access to a new source of moderately low-potential electrons (H_2_S as opposed to H_2_) that could be used together with light energy to generate reduced ferredoxin for the purpose of CO_2_ fixation via the ferredoxin-dependent acetyl-CoA pathway. The need for such function in a microorganism would be alleviated with the development of heterotrophic capacities and acquisition of additional pathways for energy production, allowing for the loss of the non-utilized chlorophyll synthesis pathway.

The second scenario posits that members of the Binatota are indeed phototrophs, possessing a complete pathway for chlorophyll biosynthesis and a novel type of reaction center that is bioinformatically unrecognizable. A minimal photosynthetic electron transport chain, similar to *Chloroflexus aurantiacus* ^101^, with the yet-unidentified reaction center, quinone, alternate complex III (or complex III) and some type of cytochrome c would possibly be functional. Under such scenario, members of the Binatota would be an extremely versatile photoheterotrophic facultative methylotrophic lineage. While such versatility, especially coupling methylotrophy to phototrophy, is rare ^102^, it has previously been observed in some Rhodospirillaceae species ^103^. A third scenario is that Binatota are capable of chlorophyll production, but still incapable of conducting photosynthesis. Under this scenario, genes missed in the pathway are due to shortcomings associated with *in-silico* prediction and conservative gene annotation. For example, the missing *bchM* (E.C.2.1.1.11) could possibly be encoded for by general methyltransferases (EC: 2.1.1.-), the missing *bciC* (EC:3.1.1.100) could possibly be encoded for by general hydrolases (EC: 3.1.1.-), while the missing *bchF* (EC:4.2.1.165) or *bchV* (EC:4.2.1.169) could possibly be encoded for by general hydratases (EC: 4.2.1.-).

Encountering incomplete pathways in genomes of uncultured lineages is an exceedingly common occurrence in SAG and MAG analysis ^104, 105^. In many cases, this could plausibly indicate an incomplete contribution to a specific biogeochemical process, e.g. incomplete denitrification of nitrate to nitrite but not ammonia ^105^, or reduction of sulfite, but not sulfate, to sulfide ^106^, provided the thermodynamic feasibility of the proposed partial pathway, and, preferably, prior precedence in pure cultures. In other cases, a pattern of absence of peripheral steps could demonstrate the capability for synthesis of a common precursor, e.g., synthesis of precorrin-2 from uroporphyrinogen, but lack of the peripheral pathway for corrin ring biosynthesis leading to an auxotrophy for vitamin B12. Such auxotrophies are common in the microbial world and could be alleviated by nutrient uptake from the outside environment ^107^ or engagement in a symbiotic lifestyle ^108^. However, arguments for metabolic interdependencies, syntrophy, or auxotrophy could not be invoked to explain the consistent absence of specific genes in a dedicated pathway, such as bacteriochlorophyll biosynthesis, especially when analyzing a large number of genomes from multiple habitats. As such, we here raise awareness that using a certain occurrence threshold to judge a pathway’s putative functionality could lead to misinterpretations of organismal metabolic capacities due to the frequent occurrence of partial, non-functional, pathways and “gene shrapnel” in microbial genomes.

In conclusion, our work provides a comprehensive assessment of the yet-uncultured phylum Binatota, and highlights its aerobic methylotrophic and alkane degradation capacities, as well as its carotenoid production, and abundance of bacteriochlorophyll synthesis genes in its genomes. We also propose a role for this lineage in mitigating methane and methanol emissions from terrestrial and freshwater ecosystems, alkanes degradation in hydrocarbon-rich habitats, and nutritional symbiosis with marine sponges. We present specific scenarios that could explain the unique pattern of chlorophyll biosynthesis gene occurrence, and stress the importance of detailed analysis of pathways completion patterns for appropriate functional assignments in genomes of uncultured taxa.

## Materials and Methods

### Genomes

All genomes classified as belonging to the Binatota in the GTDB database (n=22 MAGs, April 2020) were downloaded as assemblies from NCBI. In addition, 128 metagenome-assembled genomes with the classification “Bacteria;UBP10” were downloaded from the IMG/M database (April 2020). These genomes were recently assembled from public metagenomes as part of a wider effort to generate a genomic catalogue of Earth’s microbiome ^16^. Finally, 6 metagenome-assembled genomes were obtained as part of the Microbial Dark Matter MDM-II project. CheckM ^109^ was utilized for estimation of genome completeness, strain heterogeneity, and genome contamination. Only genomes with >70% completion and <10% contamination (n=108) were retained for further analysis (Tables S1, S2). MAGs were classified as high-, or medium-quality drafts based on the criteria set forth by ^18^.

### Phylogenetic analysis

Taxonomic classifications followed the Genome Taxonomy Database (GTDB) release r89 ^14, 110^, and were carried out using the classify_workflow in GTDB-Tk ^111^ (v1.1.0). Phylogenomic analysis utilized the concatenated alignment of a set of 120 single-copy marker genes ^14, 110^ generated by the GTDB-Tk. Maximum-likelihood phylogenomic tree was constructed in RAxML ^112^ (with a cultured representative of the phylum Deferrisomatota as the outgroup). SSU rRNA gene-based phylogenetic analysis was also conducted using 16S rRNA gene sequences extracted from genomes using RNAmmer ^113^. Putative taxonomic ranks were deduced using average amino acid identity (AAI; calculated using AAI calculator [http://enve-omics.ce.gatech.edu/]), with the arbitrary cutoffs 56%, and 68% for family, and genus, respectively.

### Annotation

Protein-coding genes in genomic bins were predicted using Prodigal ^114^. For initial prediction of function, pangenomes were constructed for each order in the phylum Binatota separately using PIRATE ^115^ with percent identity thresholds of [40, 45, 50, 55, 60, 65, 70, 75, 80, 90], a cd-hit step size of 1, and cd-hit lowest percent id of 90. The longest sequence for each PIRATE-identified allele was chosen as a representative and assembled into a pangenome. These pangenomes were utilized to gain preliminary insights on the metabolic capacities and structural features of different orders. BlastKOALA ^116^ was used to assign protein-coding genes in each of the pangenomes constructed to KEGG orthologies (KO), which were subsequently visualized using KEGG mapper ^117^. Analysis of specific capabilities and functions of interest was conducted on individual genomic bins by building and scanning hidden markov model (HMM) profiles. All predicted protein-coding genes in individual genomes were searched against custom-built HMM profiles for genes encoding C1, alkanes, and fatty acids metabolism, C1 assimilation, [NiFe] hydrogenases, electron transport chain complexes, and carotenoid and chlorophyll biosynthesis. To build the HMM profiles, Uniprot reference sequences for all genes with an assigned KO number were downloaded, aligned using Clustal-omega ^118^, and the alignment was used to build an HMM profile using hmmbuild (HMMER 3.1b2). For genes not assigned a KO number (e.g. alternative complex III genes, different classes of cytochrome c family, cytochrome P450 medium-chain alkane hydroxylase cyp153, methanol dehydrogenase MNO/MDO family), a representative protein was compared against the KEGG Genes database using Blastp and significant hits (those with e-values < e-80) were downloaded and used to build HMM profiles as explained above. The custom-built HMM profiles were then used to scan the analyzed genomes for significant hits using hmmscan (HMMER 3.1b2) with the option -T 100 to limit the results to only those profiles with an alignment score of at least 100. Further confirmation was achieved through phylogenetic assessment and tree building procedures, in which potential candidates identified by hmmscan were aligned to the reference sequences used to build the custom HMM profiles using Clustal-omega ^118^, followed by maximum likelihood phylogenetic tree construction using FastTree ^119^. Only candidates clustering with reference sequences were deemed true hits and were assigned to the corresponding KO.

### Search for photosynthetic reaction center

Identification of genes involved in chlorophyll biosynthesis in Binatota genomes prompted us to search the genomes for photosynthetic reaction center genes. HMM profiles for Reaction Center Type 1 (RC1; PsaAB), and Reaction Center Type 2 (RC2; PufLM and PsbD_1_D_2_) were obtained from the pfam database (pfam00223 and pfam00124, respectively). Additionally, HMM profiles were built for PscABCD (Chlorobia-specific), PshA/B (Heliobacteria-specific) ^120^, as well as the newly identified Psa-like genes from Chloroflexota ^121^. The HMM profiles were used to search Binatota genomes for potential hits using hmmscan. To guard against overlooking a distantly related reaction center, we relaxed our homology criteria (by not including -T or -E options during the hmmscan). An additional search using a structurally-informed reaction center alignment ^120, 122^ was also performed. The best potential hits were modeled using the SWISS-MODEL homology modeler ^123^ to check for veracity. Since the core subunits of Type 1 RC proteins are predicted to have 11 transmembrane *α*-helices ^124, 125^, while type 2 RC are known to contain five transmembrane helices ^124, 126^, we also searched for all predicted proteins harboring either 5 or 11 transmembrane domains using TMHMM ^127^. All identified 5- or 11-helix-containing protein-coding sequences were searched against GenBank protein nr database to identify and exclude all sequences with a predicted function. All remaining 5- or 11-helix-containing proteins with no predicted function were then submitted to SWISS-MODEL homology modeler using the automated mode to predict homology models.

### Classification of [NiFe] hydrogenase sequences

All sequences identified as belonging to the respiratory O_2_-tolerant H_2_-uptake [NiFe] hydrogenase large subunit (HyaA) were classified using the HydDB web tool ^128^.

### Particulate methane monooxygenase 3D model prediction and visualization

SWISS-MODEL ^123^ was used to construct pairwise sequence alignments of predicted Binatota particulate methane monooxygenase with templates from *Methylococcus capsulatus* str. Bath (pdb: 3RGB), and for predicting tertiary structure models. Predicted models were superimposed on the template enzyme in PyMol (Version 2.0 Schrödinger, LLC).

### Ecological distribution of Binatota

We queried 16S rRNA sequence databases using representative 16S rRNA gene sequences from six out of the seven Binatota orders (order UBA12015 genome assembly did not contain a 16S rRNA gene). Two databases were searched: 1. GenBank nucleotide (nt) database (accessed in July 2020) using a minimum identity threshold of 90%, *≥*80% subject length alignment for near full-length query sequences or *≥*80% query length for non-full-length query sequences, and a minimum alignment length of 100 bp, and 2. The IMG/M 16S rRNA public assembled metagenomes ^129^ using a cutoff e-value of 1e^-10^, percentage similarity *≥* 90%, and either *≥*80% subject length for full-length query sequences or *≥*80% query length for non-full-length query sequences. Hits satisfying the above criteria were further trimmed after alignment to the reference sequences from each order using Clustal-omega and inserted into maximum likelihood phylogenetic trees in FastTree (v 2.1.10, default settings). The ecological distribution for each of the Binatota orders was then deduced from the environmental sources of their hits. All environmental sources were classified according to the GOLD ecosystem classification scheme ^130^.

### Data availability

Genomic bins, predicted proteins, and extended data for Figures 2-7 and for Figures 8 and S1a are available at https://github.com/ChelseaMurphy/Binatota. Maximum likelihood trees (Figure 1 and Figure S1a) can be accessed at: https://itol.embl.de/shared/1WgxEjrQfEYWk. Maximum likelihood trees for chlorophyll biosynthesis genes are available at https://itol.embl.de/shared/34y3BUHcQd7Lh.

## Acknowledgements

This work has been supported by NSF grants 2016423 (to NHY and MSE), 1441717 and 1826734 (to RS). We thank Dr. Kevin Redding (Arizona State University) for helpful discussions. Work conducted by the U.S. Department of Energy Joint Genome Institute, a DOE Office of Science User Facility, is supported under Contract No. DE-AC02-05CH11231. J.R.S. is supported by NASA Astrobiology Rock Powered Life and was granted U.S. Forest Service permit #MLD15053 to conduct field work on Cone Pool and the Little Hot Creek, Mammoth Lakes, California. Thanks to students and participants of the 2014 – 2016 International Geobiology Course for research works on Cone Pool.

## References

1. Hug LA, et al. Critical biogeochemical functions in the subsurface are associated with bacteria from new phyla and little studied lineages. Environ. Microbiol. 18, 159–173 (2016).

2. Engelberts JP, Robbins SJ, de Goeij JM, Aranda M, Bell SC, Webster NS. Characterization of a sponge microbiome using an integrative genome-centric approach. ISME J. 14, 1100–1110 (2020).

3. Vavourakis CD, et al. Metagenomes and metatranscriptomes shed new light on the microbial-mediated sulfur cycle in a Siberian soda lake. BMC Biol. 17, 69 (2019).

4. Hu P, et al. Simulation of Deepwater Horizon oil plume reveals substrate specialization within a complex community of hydrocarbon degraders. Proc. Natl. Acad. Sci. USA 114, 7432–7437 (2017).

5. Doud DFR, et al. Function-driven single-cell genomics uncovers cellulose-degrading bacteria from the rare biosphere. ISME J 14, 659–675 (2019).

6. Anantharaman K, et al. Expanded diversity of microbial groups that shape the dissimilatory sulfur cycle. ISME J 12, 1715–1728 (2018).

7. Becraft ED, et al. Rokubacteria: Genomic giants among theuncultured bacterial phyla. Front. Micorobiol. 8, 2264 (2017).

8. Rinke R, et al. A phylogenomic and ecological analysis of the globally abundant Marine Group II archaea (Ca. Poseidoniales ord. nov.). ISME J. 13, 663–675 (2019).

9. Farag IF, Davis JP, Youssef NH, Elshahed MS. Global patterns of abundance, diversity and community structure of the Aminicenantes (Candidate Phylum OP8). PloS one 9, e92139 (2014).

10. Zhou Z, Tran PQ, Kieft K, Anantharaman K. Genome diversification in globally distributed novel marine Proteobacteria is linked to environmental adaptation. ISME J. 14, 2060–2077 (2020).

11. Youssef NH, Blainey PC, Quake SR, Elshahed MS. Partial genome assembly for a candidate division OP11 single cell from an anoxic spring (Zodletone Spring, Oklahoma). Appl. Environ. Microbiol. 77, 7804–7814 (2011).

12. Rinke C, et al. Insights into the phylogeny and coding potential of microbial dark matter. Nature 499, 431–437 (2013).

13. Beam JP, Becraft ED, Brown KM, Schulz F, Jarett JK. Ancestral absence of electron transport chains in Patescibacteria and DPANN. Front. Micorobiol. https://doi.org/10.3389/fmicb.2020.01848 (2020).

14. Parks DH, et al. A standardized bacterial taxonomy based on genome phylogeny substantially revises the tree of life. Nat. Biotechnol. 36, 996–1004 (2018)

15. Parks DH, et al. Recovery of nearly 8,000 metagenome-assembled genomes substantially expands the tree of life. Nat. Microbiol.y 2, 1533–1542 (2017).

16. Nayfach S, et al. A genomic catalogue of Earth’s microbiomes. Nat. Biotechnol. Accpeted, (2020).

17. Chuvochina M, et al. The importance of designating type material for uncultured taxa. Syst Appl Microbiol 42, 15–21 (2018).

18. Bowers RM, et al. Minimum information about a single amplified genome (MISAG) and a metagenome-assembled genome (MIMAG) of bacteria and archaea. Nat. Biotechnol. 35, 725–731 (2017).

19. Quast C, et al. The SILVA ribosomal RNA gene database project: improved data processing and web-based tools. Nucl. Acids Res. 41, D590–596 (2013).

20. Hektor HJ, Kloosterman H, Dijkhuizen L. Nicotinoprotein methanol dehydrogenase enzymes in Gram-positive methylotrophic bacteria. J. Mol. Cat. B. 8, 103–109 (2000).

21. Kalyuzhnaya MG, Hristova KR, Lidstrom ME, Chistoserdova L. Characterization of a novel methanol dehydrogenase in representatives of Burkholderiales: implications for environmental detection of methylotrophy and evidence for convergent evolution. J. Bacteriol. 190, 3817–3823 (2008).

22. Chistoserdova L. Modularity of methylotrophy, revisited. Environ. Microbiol. 13, 2603–2622 (2011).

23. Erikstad HA, Jensen S, Keen TJ, Birkeland NK. Differential expression of particulate methane monooxygenase genes in the verrucomicrobial methanotroph *’Methylacidiphilum kamchatkense*’ Kam1. Extremophiles 16, 405–409 (2012).

24. Ettwig KF, et al. Nitrite-driven anaerobic methane oxidation by oxygenic bacteria. Nature 464, 543–548 (2010).

25. Iguchi H, Yurimoto H, Sakai Y. Soluble and particulate methane monooxygenase gene clusters of the type I methanotroph *Methylovulum miyakonense* HT12. FEMS Microbiol. Lett. 312, 71–76 (2010).

26. Ricke P, Erkel C, Kube M, Reinhardt R, Liesack W. Comparative analysis of the conventional and novel pmo (particulate methane monooxygenase) operons from *Methylocystis* strain SC2. Appl. Environ. Microbiol. 70, 3055–3063 (2004).

27. Fisher OS, et al. Characterization of a long overlooked copper protein from methane- and ammonia-oxidizing bacteria. Nat. Commun. 9, 4276 (2018).

28. Rochman FF, et al. Novel copper-containing membrane monooxygenases (CuMMOs) encoded by alkane-utilizing Betaproteobacteria. ISME J. 14, 714–726 (2020).

29. Knief C. Diversity and habitat preferences of cultivated and uncultivated aerobic methanotrophic bacteria evaluated based on pmoA as molecular marker. Front. Microbiol. 6, 1346 (2015).

30. Li M, Jain S, Baker BJ, Taylor C, Dick GJ. Novel hydrocarbon monooxygenase genes in the metatranscriptome of a natural deep-sea hydrocarbon plume. Environ. Microbiol. 16, 60–71 (2014).

31. Sheik CS, Jain S, Dick GJ. Metabolic flexibility of enigmatic SAR324 revealed through metagenomics and metatranscriptomics. Environ. Microbiol. 16, 304–317 (2014).

32. Kalyuzhnaya MG, Zabinsky R, Bowerman S, Baker DR, Lidstrom ME, Chistoserdova L. Fluorescence in situ hybridization-flow cytometry-cell sorting-based method for separation and enrichment of type I and type II methanotroph populations. Appl. Environ. Microbiol. 72, 4293–4301 (2006).

33. Hamamura N, Yeager CM, Arp DJ. Two distinct monooxygenases for alkane oxidation in Nocardioides sp. strain CF8. Appl. Environ. Microbiol. 67, 4992–4998 (2001).

34. Lessmeier L, Hoefener M, Wendisch VF. Formaldehyde degradation in Corynebacterium glutamicum involves acetaldehyde dehydrogenase and mycothiol-dependent formaldehyde dehydrogenase. Microbiology (Reading, England) 159, 2651–2662 (2013).

35. Dubey AA, Wani SR, Jain V. Methylotrophy in Mycobacteria: dissection of the methanol metabolism pathway in *Mycobacterium smegmatis*. J. Bacteriol. 200, e00288–18 (2018).

36. Kornberg HL, Krebs HA. Synthesis of cell constituents from C2-units by a modified tricarboxylic acid cycle. Nature 179, 988–991 (1957).

37. Alber BE, Spanheimer R, Ebenau-Jehle C, Fuchs G. Study of an alternate glyoxylate cycle for acetate assimilation by *Rhodobacter sphaeroides*. Mol. Microbiol. 61, 297–309 (2006).

38. Chen Q, Janssen DB, Witholt B. Growth on octane alters the membrane lipid fatty acids of Pseudomonas oleovorans due to the induction of alkB and synthesis of octanol. J. Bacteriol. 177, 6894–6901 (1995).

39. van Beilen JB, Funhoff EG. Alkane hydroxylases involved in microbial alkane degradation. Appl. Microbiol. Biotechnol. 74, 13–21 (2007).

40. Li L, et al. Crystal structure of long-chain alkane monooxygenase (LadA) in complex with coenzyme FMN: unveiling the long-chain alkane hydroxylase. J. Mol. Biol. 376, 453–465 (2008).

41. Nagata Y, Miyauchi K, Damborsky J, Manova K, Ansorgova A, Takagi M. Purification and characterization of a haloalkane dehalogenase of a new substrate class from a gamma-hexachlorocyclohexane-degrading bacterium, *Sphingomonas paucimobilis* UT26. Appl. Environ. Microbiol. 63, 3707–3710 (1997).

42. Sun C, et al. Structure of the alternative complex III in a supercomplex with cytochrome oxidase. Nature 557, 123–126 (2018).

43. Choi DW, et al. The membrane-associated methane monooxygenase (pMMO) and pMMO-NADH:quinone oxidoreductase complex from *Methylococcus capsulatus* Bath. J. Bacteriol.y 185, 5755–5764 (2003).

44. Nguyen HH, Elliott SJ, Yip JH, Chan SI. The particulate methane monooxygenase from *Methylococcus capsulatus* (Bath) is a novel copper-containing three-subunit enzyme. Isolation and characterization. J. Biol. Chem. 273, 7957–7966 (1998).

45. Wulff P, Day CC, Sargent F, Armstrong FA. How oxygen reacts with oxygen-tolerant USA 111, 6606–6611 (2014).

46. Sargent F. The Model [NiFe]-Hydrogenases of *Escherichia coli*. Adv. Microb. Physiol. 68, 433–507 (2016).

47. Volbeda A, Darnault C, Parkin A, Sargent F, Armstrong FA, Fontecilla-Camps JC. Crystal structure of the O(2)-tolerant membrane-bound hydrogenase 1 from *Escherichia coli* in complex with its cognate cytochrome b. Structure 21, 184–190 (2013).

48. Carere CR, et al. Mixotrophy drives niche expansion of verrucomicrobial methanotrophs. ISME J. 11, 2599–2610 (2017).

49. Chistoserdova L, Kalyuzhnaya MG, Lidstrom ME. The expanding world of methylotrophic metabolism. Ann. Rev. Microbiol. 63, 477–499 (2009).

50. Boden R, Thomas E, Savani P, Kelly DP, Wood AP. Novel methylotrophic bacteria isolated from the River Thames (London, UK). Environ. Microbiol. 10, 3225–3236 (2008).

51. McTaggart TL, et al. Genomics of methylotrophy in Gram-positive methylamine-utilizing bacteria. Microorganisms 3, 94–112 (2015).

52. Pol A, Heijmans K, Harhangi HR, Tedesco D, Jetten MSM, Op den Camp HJM. Methanotrophy below pH 1 by a new Verrucomicrobia species. Nature 450, 874–878 (2007).

53. Ettwig KF, van Alen T, van de Pas-Schoonen KT, Jetten MS, Strous M. Enrichment and molecular detection of denitrifying methanotrophic bacteria of the NC10 phylum. Appl. Environ. Microbiol. 75, 3656–3662 (2009).

54. Diamond S, et al. Mediterranean grassland soil C-N compound turnover is dependent on rainfall and depth, and is mediated by genomically divergent microorganisms. Nat. Microbiol. 4, 1356–1367 (2019).

55. Butterfield CN, et al. Proteogenomic analyses indicate bacterial methylotrophy and archaeal heterotrophy are prevalent below the grass root zone. PeerJ 4, e2687 (2016).

56. Davamani V, Parameswari E, Arulmani S. Mitigation of methane gas emissions in flooded paddy soil through the utilization of methanotrophs. Sci Tot. Environ. 726, 138570 (2020).

57. Khmelenina VN, Colin Murrell J, Smith TJ, Trotsenko YA. Physiology and biochemistry of the aerobic methanotrophs. In: Aerobic utilization of hydrocarbons, oils and lipids (ed Rojo F). Springer International Publishing (2018).

58. Mohammadi SS, Pol A, van Alen T, Jetten MSM, Op den Camp HJM. Ammonia oxidation and nitrite reduction in the verrucomicrobial methanotroph *Methylacidiphilum fumariolicum* SolV. Front. Microbiol. 8, 1901 (2017).

59. Prince RC, Amande TJ, McGenity TJ. Prokaryotic hydrocarbon degraders. In: Taxonomy, genomics and ecophysiology of hydrocarbon-degrading microbes (ed McGenity TJ). Springer International Publishing (2019).

60. Le Mer J, Roger P. Production, oxidation, emission and consumption of methane by soils: A review. Eur. J. Soil Biol. 37, 25–50 (2001).

61. Wang Z, Zeng D, Patrick WH. Methane emissions from natural wetlands. Environ. Monit. Assess.42, 143–161 (1996).

62. He S, et al. Patterns in wetland microbial community composition and functional gene repertoire associated with methane emissions. mBio 6, e00066–00015 (2015).

63. Angle JC, et al. Methanogenesis in oxygenated soils is a substantial fraction of wetland methane emissions. Nat. Commun. 8, 1567 (2017).

64. Kolb S. Aerobic methanol-oxidizing Bacteria in soil. FEMS Microbiol. Lett. 300, 1–10 (2009).

65. Conrad R. The global methane cycle: recent advances in understanding the microbial processes involved. Environ. Microbiol. Rep. 1, 285–292 (2009).

66. McCollom TM. Laboratory simulations of abiotic hydrocarbon formation in earth’s deep subsurface. Rev.n Mineral. Geochem. 75, 467–494 (2013).

67. Wang W, Li Z, Zeng L, Dong C, Shao Z. The oxidation of hydrocarbons by diverse heterotrophic and mixotrophic bacteria that inhabit deep-sea hydrothermal ecosystems. ISME J. 14, 1994–2006 (2020).

68. Pasche N, et al. Methane sources and sinks in Lake Kivu. J. Geophys. Res. Biogeosci. 116, G03006 (2011).

69. Borges AV, Abril G, Delille B, Descy J-P, Darchambeau F. Diffusive methane emissions to the atmosphere from Lake Kivu (Eastern Africa). J. Geophys. Res. Biogeosci. 116, G03032 (2011).

70. Llirós M, et al. Microbial Ecology of Lake Kivu. In: Lake Kivu: Limnology and biogeochemistry of a tropical great lake (eds Descy J-P, Darchambeau F, Schmid M). Springer Netherlands (2012).

71. Chistoserdova L. Methylotrophy in a lake: from metagenomics to single-organism physiology. Appl. Environ. Microbiol. 77, 4705–4711 (2011).

72. Chistoserdova L. The distribution and evolution of c1 transfer enzymes and evolution of the Planctomycetes. In: Planctomycetes: Cell Structure, Origins and Biology (ed Fuerst JA). Humana Press (2013).

73. Auman AJ, Lidstrom ME. Analysis of sMMO-containing type I methanotrophs in Lake Washington sediment. Environ. Microbiol. 4, 517–524 (2002).

74. Auman AJ, Stolyar S, Costello AM, Lidstrom ME. Molecular characterization of methanotrophic isolates from freshwater lake sediment. Appl. Environ. Microbiol. 66, 5259–5266 (2000).

75. Chistoserdova L. Methylotrophs in natural habitats: current insights through metagenomics. Appl. Microbiol. Biotechnol. 99, 5763–5779 (2015).

76. Kuivila KM, Murray JW, Devol AH, Lidstrom ME, Reimers CE. Methane cycling in the sediments of Lake Washington. Limnol. Oceanogr. 33, 571–581 (1988).

77. Arellano SM, et al. Deep sequencing of *Myxilla* (*Ectyomyxilla*) *methanophila*, an epibiotic sponge on cold-seep tubeworms, reveals methylotrophic, thiotrophic, and putative hydrocarbon-degrading microbial associations. Microb. Ecol. 65, 450–461 (2013).

78. Tian RM, Zhang W, Cai L, Wong YH, Ding W, Qian PY. Genome reduction and microbe-host interactions drive adaptation of a sulfur-oxidizing bacterium associated with a cold seep sponge. mSystems 2, (2017).

79. Rubin-Blum M, et al. Fueled by methane: deep-sea sponges from asphalt seeps gain their nutrition from methane-oxidizing symbionts. ISME J. 13, 1209–1225 (2019).

80. Hashimoto H, Uragami C, Cogdell RJ. Carotenoids and photosynthesis. Subcell. Biochem. 79, 111–139 (2016).

81. Saidi-Mehrabad A, et al. *Methylicorpusculum oleiharenae* gen. nov., sp. nov., an aerobic methanotroph isolated from an oil sands tailings pond. Int. J. Syst. Evol. Microbiol. 70, 2499–2508 (2020).

82. Wang FQ, et al. *Carboxylicivirga sediminis* sp. nov., isolated from coastal sediment. Int. J. Syst. Evol. Microbiol. 68, 1896–1901 (2018).

83. Asker D, Awad TS, Beppu T, Ueda K. *Deinococcus misasensis* and *Deinococcus roseus*, novel members of the genus *Deinococcus*, isolated from a radioactive site in Japan. Syst. Appl. Microbiol. 31, 43–49 (2008).

84. Zhou EM, et al. *Thermus sediminis* sp. nov., a thiosulfate-oxidizing and arsenate-reducing organism isolated from Little Hot Creek in the Long Valley Caldera, California. Extremophiles 22, 983–991 (2018).

85. Sanford RA, Cole JR, Tiedje JM. Characterization and description of *Anaeromyxobacter dehalogenans* gen. nov., sp. nov., an aryl-halorespiring facultative anaerobic myxobacterium. Appl. Environ. Microbiol. 68, 893–900 (2002).

86. Fariq A, Yasmin A, Jamil M. Production, characterization and antimicrobial activities of bio-pigments by *Aquisalibacillus elongatus* MB592, *Salinicoccus sesuvii* MB597, and *Halomonas aquamarina* MB598 isolated from Khewra Salt Range, Pakistan. Extremophiles 23, 435–449 (2019).

87. Ungers GE, Cooney JJ. Isolation and characterization of carotenoid pigments of *Micrococcus roseus*. J. Bacteriol. 96, 234–241 (1968).

88. Bondoso J, Albuquerque L, Nobre MF, Lobo-da-Cunha A, da Costa MS, Lage OM. *Roseimaritima ulvae* gen. nov., sp. nov. and *Rubripirellula obstinata* gen. nov., sp. nov. two novel planctomycetes isolated from the epiphytic community of macroalgae. Syst. Appl. Microbiol. 38, 8–15 (2015).

89. Chen S, Sun S, Xu Y, Liu HC. *Halococcus salsus* sp. nov., a novel halophilic archaeon isolated from rock salt. Int. J. Syst. Evol. Microbiol. 68, 3754–3759 (2018).

90. Grogan DW. Phenotypic characterization of the archaebacterial genus *Sulfolobus*: comparison of five wild-type strains. J. Bacteriol. 171, 6710–6719 (1989).

91. Fiedor J, Sulikowska A, Orzechowska A, Fiedor L, Burda K. Antioxidant effects of carotenoids in a model pigment-protein complex. Acta Biochim. Polon. 59, 61–64 (2012).

92. Krisko A, Radman M. Biology of extreme radiation resistance: the way of *Deinococcus radiodurans*. *Cold Spring Harbor Pers*. Biol. 5, a012765 (2013).

93. Du X-j, Wang X-y, Dong X, Li P, Wang S. Characterization of the desiccation tolerance of *Cronobacter sakazakii* strains. Front Microbiol. 9, 2867 (2018).

94. Bowman JP, Sly LI, Nichols PD, Hayward AC. Revised taxonomy of the methanotrophs: Description of *Methylobacter* gen. nov., Emendation of *Methylococcus*, Validation of *Methylosinus* and *Methylocystis* Species, and a proposal that the family Methylococcaceae includes only the group I methanotrophs. Int. J. Syst. Evol. Microbiol. 43, 735–753 (1993).

95. Irvine IC, Brigham CA, Suding KN, Martiny JB. The abundance of pink-pigmented facultative methylotrophs in the root zone of plant species in invaded coastal sage scrub habitat. PloS one 7, e31026 (2012).

96. Mendler K, Chen H, Parks DH, Lobb B, Hug LA, Doxey AC. AnnoTree: visualization and exploration of a functionally annotated microbial tree of life. Nucl. Acids Res. 47, 4442–4448 (2019).

97. Meng J, et al. An uncultivated crenarchaeota contains functional bacteriochlorophyll a synthase. ISME J. 3, 106–116 (2009).

98. Liu R, Cai R, Zhang J, Sun C. Heimdallarchaeota harness light energy through photosynthesis. bioRxiv, 2020.2002.2020.957134 (2020).

99. Martin WF, Bryant DA, Beatty JT. A physiological perspective on the origin and evolution of photosynthesis. FEMS Mmicrobiol. Rev. 42, 205–231 (2018).

100. White SN, Chave AD, Reynolds GT, Van Dover CL. Ambient light emission from hydrothermal vents on the Mid-Atlantic Ridge. Geophys. Res. Lett. 29, 34–31-34-34 (2002).

101. Gao X, Xin Y, Bell PD, Wen J, Blankenship RE. Structural analysis of alternative complex III in the photosynthetic electron transfer chain of *Chloroflexus aurantiacus*. Biochemistry 49, 6670–6679 (2010).

102. Chistoserdova L, Lidstrom ME. Aerobic methylotrophic prokaryotes. In: The Prokaryotes: prokaryotic physiology and biochemistry (eds Rosenberg E, DeLong EF, Lory S, Stackebrandt E, Thompson F). Springer Berlin Heidelberg (2013).

103. Quayle JR, Pfennig N. Utilization of methanol by rhodospirillaceae. Arch. Microbiol. 102, 193–198 (1975).

104. Anantharaman K, et al. Thousands of microbial genomes shed light on interconnected biogeochemical processes in an aquifer system. Nat. Commun. 7, 13219 (2016).

105. Hug LA, Co R. It takes a village: microbial communities thrive through interactions and metabolic handoffs. mSystems 3, e00152–00117 (2018).

106. Colman DR, Lindsay MR, Amenabar MJ, Fernandes-Martins MC, Roden ER, Boyd ES. Phylogenomic analysis of novel Diaforarchaea is consistent with sulfite but not sulfate reduction in volcanic environments on early Earth. ISME J. 14, 1316–1331 (2020).

107. Garcia SL, Buck M, McMahon KD, Grossart H-P, Eiler A, Warnecke F. Auxotrophy and intrapopulation complementary in the ‘interactome’ of a cultivated freshwater model community. Mol. Ecol. 24, 4449–4459 (2015).

108. Croft MT, Lawrence AD, Raux-Deery E, Warren MJ, Smith AG. Algae acquire vitamin B12 through a symbiotic relationship with bacteria. Nature 438, 90–93 (2005).

109. Parks DH, Imelfort M, Skennerton CT, Hugenholtz P, Tyson GW. CheckM: assessing the quality of microbial genomes recovered from isolates, single cells, and metagenomes. Genome Res. 25, 1043–1055 (2015).

110. Parks DH, Chuvochina M, Chaumeil PA, Rinke C, Mussig AJ, Hugenholtz P. A complete domain-to-species taxonomy for Bacteria and Archaea. Nat. Biotechnol. 38,1079–1086 (2020).

111. Chaumeil PA, Mussig AJ, Hugenholtz P, Parks DH. GTDB-Tk: a toolkit to classify genomes with the Genome Taxonomy Database. Bioinformatics 36, 1925–1927 (2019).

112. Stamatakis A. RAxML version 8: a tool for phylogenetic analysis and post-analysis of large phylogenies. Bioinformatics 30, 1312–1313 (2014).

113. Lagesen K, Hallin P, Rødland EA, Staerfeldt HH, Rognes T, Ussery DW. RNAmmer: consistent and rapid annotation of ribosomal RNA genes. Nucl. Acids Res. 35, 3100–3108 (2007).

114. Hyatt D, Chen GL, Locascio PF, Land ML, Larimer FW, Hauser LJ. Prodigal: prokaryotic gene recognition and translation initiation site identification. BMC Bioinformatics 11, 119 (2010).

115. Bayliss SC, Thorpe HA, Coyle NM, Sheppard SK, Feil EJ. PIRATE: A fast and scalable pangenomics toolbox for clustering diverged orthologues in bacteria. GigaScience 8, giz119 (2019).

116. Kanehisa M, Sato Y, Morishima K. BlastKOALA and GhostKOALA: KEGG Tools for Functional Characterization of Genome and Metagenome Sequences. J. Mol. Biol. 428, 726–731 (2016).

117. Kanehisa M, Sato Y. KEGG Mapper for inferring cellular functions from protein sequences. Prot. Sci. 29, 28–35 (2020).

118. Sievers F, Higgins DG. Clustal Omega for making accurate alignments of many protein sequences. Prot. Sci. 27, 135–145 (2018).

119. Price MN, Dehal PS, Arkin AP. FastTree 2--approximately maximum-likelihood trees for large alignments. PloS one 5, e9490 (2010).

120. Orf GS, Gisriel C, Redding KE. Evolution of photosynthetic reaction centers: insights from the structure of the heliobacterial reaction center. Photosyn. Res. 138, 11–37 (2018).

121. Tsuji J, et al. Anoxygenic phototrophic Chloroflexota member uses a Type I reaction center. bioRxiv, 2020.2007.2007.190934 (2020).

122. Sadekar S, Raymond J, Blankenship RE. Conservation of distantly related membrane proteins: photosynthetic reaction centers share a common structural core. Mol. Biol. Evol. 23, 2001–2007 (2006).

123. Waterhouse A, et al. SWISS-MODEL: homology modelling of protein structures and complexes. Nucl. Acids Res. 46, W296–w303 (2018).

124. Hohmann-Marriott MF, Blankenship RE. Evolution of photosynthesis. Ann. Rev. Plant Biol. 62, 515–548 (2011).

125. Krauß N. Structure and function of cyanobacterial photosystem I. In: Photosynthetic Protein Complexes (ed Fromme P). Wiley-Blackwell (2008).

126. Allen JP, Williams JC. Reaction centers from purple bacteria. In: Photosynthetic Protein Complexes (ed Fromme P). Wiley-Blackwell (2008).

127. Krogh A, Larsson B, von Heijne G, Sonnhammer EL. Predicting transmembrane protein topology with a hidden Markov model: application to complete genomes. J. Mol. Biol. 305, 567–580 (2001).

128. Søndergaard D, Pedersen CN, Greening C. HydDB: A web tool for hydrogenase classification and analysis. Sci. Rep. 6, 34212 (2016).

129. Chen IA, et al. IMG/M v.5.0: an integrated data management and comparative analysis system for microbial genomes and microbiomes. Nucl. Acids Res. 47, D666–d677 (2019).

130. Mukherjee S, et al. Genomes OnLine database (GOLD) v.7: updates and new features. Nucl. Acids Res. 47, D649–d659 (2019).

131. Kits KD, Klotz MG, Stein LY. Methane oxidation coupled to nitrate reduction under hypoxia by the Gammaproteobacterium *Methylomonas denitrificans*, sp. nov. type strain FJG1. Environ. Microbiol. 17, 3219–3232 (2015).

132. Tavormina PL, Orphan VJ, Kalyuzhnaya MG, Jetten MS, Klotz MG. A novel family of functional operons encoding methane/ammonia monooxygenase-related proteins in gammaproteobacterial methanotrophs. Environ. Microbiol. Rep. 3, 91–100 (2011).

133. Vorobev A, et al. Genomic and transcriptomic analyses of the facultative methanotroph *Methylocystis* sp. strain SB2 grown on methane or ethanol. Appl. Environ. Microbiol. 80, 3044–3052 (2014).

