## Supplemental document for "Methylotrophy, alkane-degradation, and pigment production as defining features of the globally distributed yet-uncultured phylum Binatota"

**Table S1.** Binatota genomes used in this study^#^, their GTDB classification and the corresponding classification in Silva and RDP databases, and the source from which they were obtained.

**Table S2.** Sequencing statistics for the genomic bins used in this study

**Table S3.** General genomic features of the studied genomes.

**Figure S1.** (A) Maximum likelihood phylogenetic tree based on the 16S rRNA gene representatives from six Binatota orders with representative hit sequences (number of sequences in parentheses following the order name) from the IMG and NCBI nt databases identified by Blastn. Orders are color coded following the color scheme in Figure 1, and the number of hits from each database are shown in parentheses. Bootstrap value (from 100 bootstraps) are shown for branches with >70% support.

(B-G) Ecological distribution of Binatota-affiliated 16S rRNA sequences. Representative 16S rRNA gene sequences from six out of the seven Binatota orders (order UBA12015 genome assembly did not contain a16S rRNA gene) were searched against Integrated Microbial Genomes & Microbiomes (IMG/M) 16S rRNA public assembled metagenomes database using Blastn and the criteria specified in Materials and Methods. Binatota orders are shown on the X-axis, while percentage abundance in different environments (classified based on the GOLD ecosystem classification scheme) are shown on the Y-axis (B). Further sub-classifications for each environment are shown for (C) terrestrial, (D) freshwater, (E) marine, (F) host-associated, and (G) engineered environments. Details including GenBank accession number of hit sequences are shown in Extended data 3.

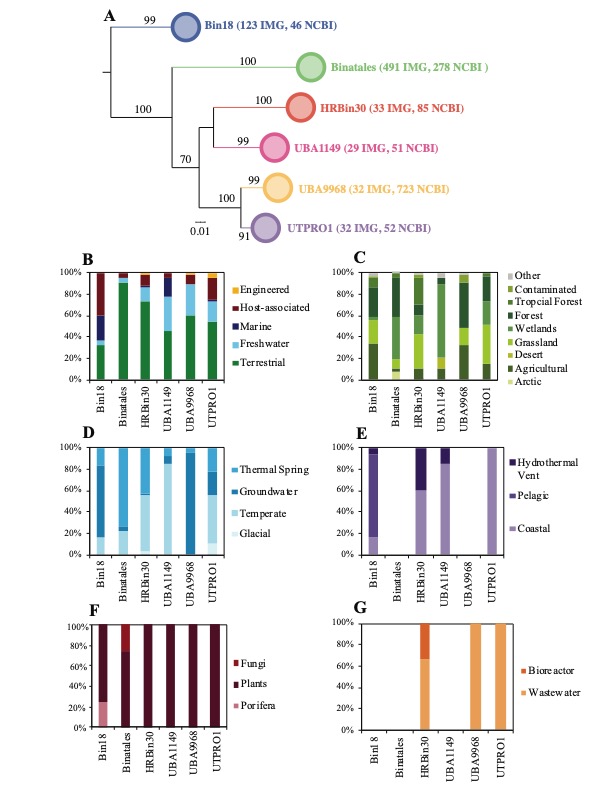

**Extended data 1.** Different sheets correspond to the heatmaps in Figures 2 through 7 in the main text. Each sheet references the protein IDs in each of the 108 Binatota genomes corresponding to the gene in the column header. The actual amino acid sequences can be found in the amino acid fasta files available at <https://github.com/ChelseaMurphy/Binatota> by searching by the protein ID in the corresponding genome file.

**Extended data 2.** Different sheets correspond to different Binatota orders. Each sheet lists the accession numbers, source, and ecosystem-level classification of the NCBI nt database hits identified using Blastn. These data were used to construct Figure 8.

**Extended data 3.** Different sheets correspond to different Binatota orders. Each sheet lists the accession numbers, source, and ecosystem-level classification of the IMG database hits identified using Blastn. These data were used to construct Figure S1.

**References**

1.	Sorensen JW, Dunivin TK, Tobin TC, Shade A. Ecological selection for small microbial genomes along a temperate-to-thermal soil gradient. *Nat. Microbiol.***4**, 55-61 (2019).

2.	Slaby BM, Hackl T, Horn H, Bayer K, Hentschel U. Metagenomic binning of a marine sponge microbiome reveals unity in defense but metabolic specialization. *ISME J.* **11**, 2465-2478 (2017).

3.	Hausmann B*, et al.* Peatland Acidobacteria with a dissimilatory sulfur metabolism. *ISME J.* **12**, 1729-1742 (2018).

4.	Parks DH*, et al.* Recovery of nearly 8,000 metagenome-assembled genomes substantially expands the tree of life. *Nat. Microbiol.* **2**, 1533-1542 (2017).

5.	Thomas SC*, et al.* Position-specific metabolic probing and metagenomics of microbial communities reveal conserved central carbon metabolic network activities at high temperatures. *Front. Microbiol.* **10**, 1427 (2019).

6.	Kato S*, et al.* Long-term cultivation and metagenomics reveal ecophysiology of previously uncultivated thermophiles involved in biogeochemical nitrogen cycle. *Microbes Environ.***33**, 107-110 (2018).

7.	Anantharaman K, Duhaime MB, Breier JA, Wendt KA, Toner BM, Dick GJ. Sulfur oxidation genes in diverse deep-sea viruses. *Science* **344**, 757-760 (2014).

8.	Anantharaman K*, et al.* Thousands of microbial genomes shed light on interconnected biogeochemical processes in an aquifer system. *Nat. Commun.* **7**, 13219 (2016).

9.	Parks DH*, et al.* A standardized bacterial taxonomy based on genome phylogeny substantially revises the tree of life. *Nat. Biotechnol.* **36**, 996–1004 (2018).

10.	Lawson CE*, et al.* Metabolic network analysis reveals microbial community interactions in anammox granules. *Nat. Commun.* **8**, 15416 (2017).
